# Selenoprotein S plays a role in translation and membrane protein biogenesis

**DOI:** 10.64898/2026.06.08.725543

**Authors:** Farid Ghelichkhani, Mariia A. Kapitonova, Atinuke Odunsi, Erfan Rahmani, Omar G. Rosas Bringas, Mohammed Hanzala Kaniyar, John LaCava, Sharon Rozovsky

**Affiliations:** Department of Chemistry and Biochemistry, University of Delaware, Newark, DE, 19716 USA; Department of Biomedical Engineering, University of Delaware, Newark, DE, 19713 USA; Laboratory of Cellular and Structural Biology, The Rockefeller University, New York City, NY, USA

**Author notes:** Corresponding author: Sharon Rozovsky, Department of Chemistry and Biochemistry, University of Delaware, Newark, DE, 19716 USA.

## Abstract

Human selenoprotein S (selenos) is part of the integrated cellular stress response and linked to protein quality control and signaling pathways. Consequently, genetic polymorphisms of selenos are associated with increased risk for diabetes, dyslipidemia, and cardiovascular diseases. Determining the specific roles of selenos in these cellular pathways and diseases has been challenging, as selenos associates with a wide range of protein complexes. Thus, to map the cellular functions of selenos and uncover their interconnections, we used affinity purification and *in vivo* crosslinking to stabilize transient protein interactions, followed by proteomics to record the resulting selenos interactome. Through mapping of selenos protein partners, we found evidence that selenos associates with complexes responsible for the insertion of membrane proteins into the ER bilayer and their connected quality control components. Furthermore, selenos is also part of metabolic, trafficking, and mitochondrial pathways. Notably, proteins involved in translation preferentially associate with selenos when its C-terminal intrinsically disordered segment containing the redox-active motif is accessible. Together, these results identify the C-terminal redox loop of selenos as a central interaction hub connecting translation with ER membrane protein biogenesis and quality control.

## Introduction

Selenoprotein S (selenos^1^, previously known as VIMP, SELS, and SEPS) is one of the 25 human selenoproteins that are distinguished by containing the reactive amino acid selenocysteine (Sec)^2^. Sec is used by nature to expand the chemical versatility of enzymes beyond the limits set by the 20 canonical amino acids^3,4^. Reaping the benefits of the rare and reactive selenium, however, comes at a high cost to the cell’s resources^5^. Thus, Sec is primarily employed as a catalytic residue in enzymes^4,6^. In humans, many selenoproteins are involved in managing reactive oxygen species and resolving cellular stress. Consistent with this pattern, selenos participates in the integrated stress response which is involved in processes responsible for protein quality control, inflammation, signaling, and differentiation^7–9^. The extensive literature, reviewed in reference^9^, implicates selenos in multiple pathways. However, its specific roles in cellular homeostasis and during cellular stress remain under active investigation.

Selenos is a positively charged, 189-amino-acid, single-pass^10^ membrane protein present in all human cells^2^, and was shown to reside primarily in the endoplasmic reticulum (ER) membrane, but also in the perinuclear membrane, and the Golgi^11^. It is a component of the ER-associated degradation (ERAD) pathway^12^ which extracts misfolded proteins from the ER or the cytosol and unfolds them for degradation by proteasome^13,14^. Selenos contributes to lipid synthesis, triglyceride storage, and adipogenesis^15–17^ and is thus linked to metabolic disorders such as obesity, type 2 diabetes mellitus, and nonalcoholic liver disease^17,18^. It also influences the NF-ĸB, Akt/mTOR, and STAT3 signaling pathways, cytokine levels, and calcium homeostasis^19,20^. Furthermore, it has been linked to several cancers^21^. While we have shown that selenos is a powerful reductase *in vitro*^22,23^, its *in vivo* substrates remain unidentified. For a protein implicated in so many processes and human diseases (reviewed in^9^), it is surprising that its knockout in cells and mice is well tolerated and in mice causes only deficiencies in calcium signaling and muscle function^8,17,24^. Despite all these insights, the following central questions remain unanswered: What does selenos do at the molecular level? Which proteins does it bind to directly? What protein complexes or supercomplexes does it form? Why do selenos’s functions require selenocysteine (Sec, U)?

Furthermore, in selenos, Sec is deployed in an unconventional manner. Many human selenoproteins have a thioredoxin fold, and their Sec is coupled to a cysteine two to three amino acids away. However, AlphaFold3^25^ predicts that selenos has three α-helices, which are followed by an extended intrinsically disordered segment (IDS) harboring the reactive Sec (Fig. 1A). This is notable because enzymes that are disordered are extremely rare. Equally unique is selenos’s C-terminal “stapled” loop, which is 15 amino acids long and formed by a selenylsulfide bond between the Sec and the resolving Cys at position 174. The intramolecular selenylsulfide bond controls the conformation of this loop (C_174_SWRPGRRGPSSGGU_188_G) and only reduced selenos will expose the Sec, enabling it to react with substrates. Due to the high reactivity of Sec, the selenylsulfide bond reforms rapidly, restoring the “stapled” loop conformation much faster than a disulfide bond could form^22,23^. Furthermore, the low redox potential of the selenylsulfide bond offers very tight control, as only a few cellular enzymes are capable of breaking it. The loop itself is rich in prolines, arginines, and glycines (Fig. 1A). Due to the prolines, the redox loop is likely to be structured and not ‘unwind’ when the selenylsulfide bond is reduced. However, structural predictions hint that the loop tryptophan may be cascading a conformational change and thus potentially changes in the loop interactions with protein partners.

**Figure 1.**
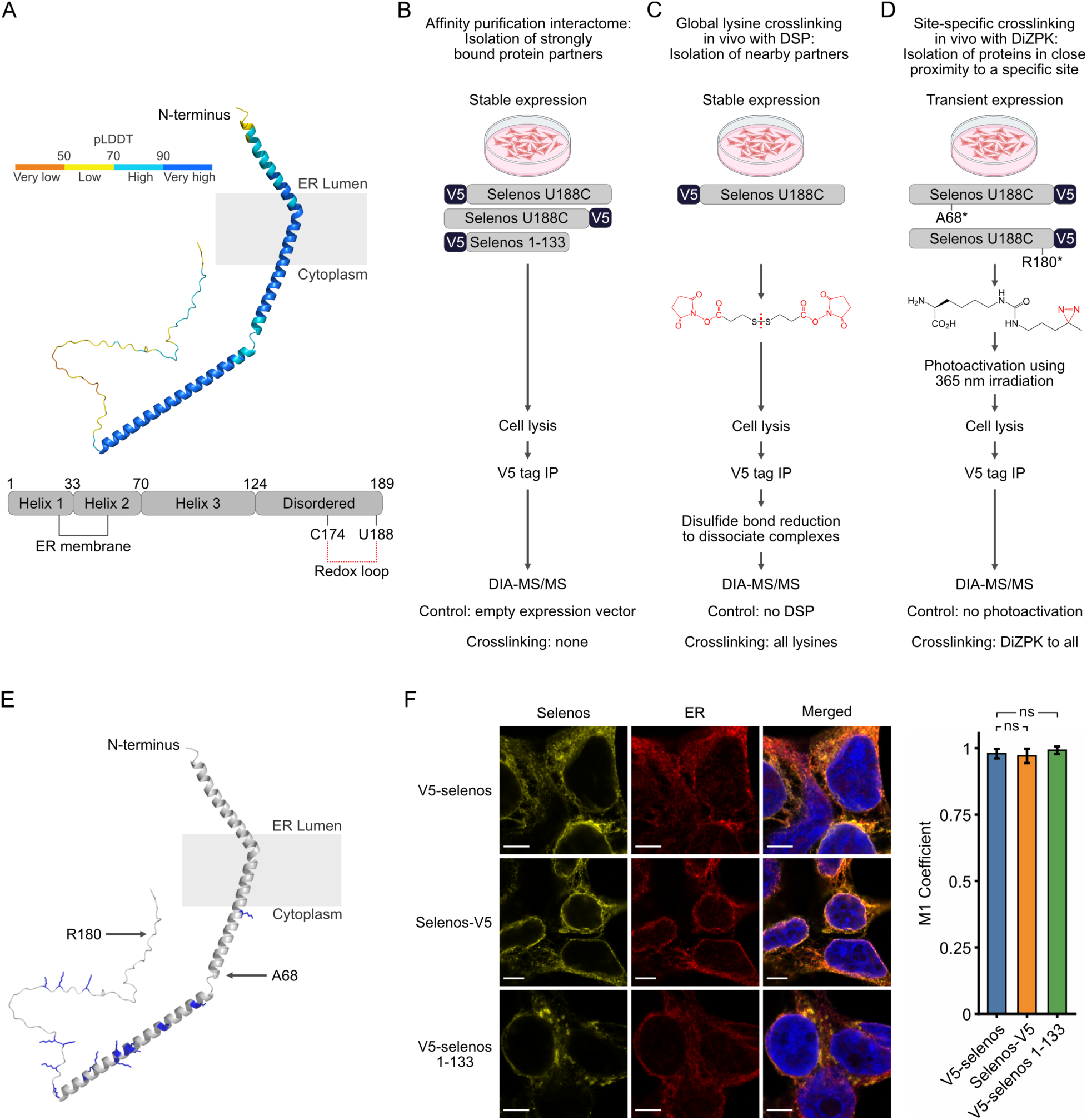
Selenos’s structural features and the experimental workflow used to identify protein partners through comparing information from affinity purifications using different selenos segments and *in vivo* crosslinking and segmentation of selenos. A) AlphaFold3^31^ prediction of selenos’s structure. Prediction confidence is color-coded by predicted local distance difference (pLDDT). The predicted transmembrane segment was identified using TMHMM. B-D) Proteomics workflows. B) Procedure for affinity purification sample preparation (NC-interactome). Flp-In T-REx 293 cells stably expressing selenos variants were lysed, and bait proteins were immunoprecipitated with V5 nanobody. Interactomes were analyzed by data-independent acquisition (DIA) mass spectrometry using five replicates. C) Procedure for global lysine crosslinking conditions (LC-interactome) using the crosslinker DSP. Flp-In T-REx 293 cells expressing V5-selenos U188C were treated with the crosslinker DSP to link proximal lysines *in vivo*. Following incubation, the cells were lysed and processed similarly to the workflow presented in B, but with an additional step to reduce the disulfide bond in DSP’s linker to disconnect crosslinked proteins after immunoprecipitation and allow their efficient digestion. Experiments were carried out in triplicates. D) Procedure for site-specific crosslinking conditions (SC-interactome) using the crosslinker DiZPK. HEK293T cells were transiently transfected with selenos-V5 with an amber codon at position 68 or 180, together with an orthogonal tRNA/aminoacyl-tRNA synthetase. The crosslinker DiZPK was supplied. The C-terminal V5 tag enabled immunoprecipitation of only DiZPK-incorporated selenos. Cultures were UV-irradiated (365 nm) prior to immunoprecipitation to induce site-specific crosslinking between selenos and nearby proteins *in vivo*. Experiments were carried out in triplicates. E) Distribution of lysine residues (blue) on the selenos structural model predicted using AlphaFold3^31^. Positions used for DiZPK incorporation are indicated by arrows. F) Immunofluorescence microscopy of selenos variants (V5 antibody) (yellow), the ER marker using calnexin antibody (red), and nuclei (stained with DAPI) (blue). Error bars show mean ± SD. The scale bar is 5 μm. The M1 coefficient quantifies selenos localization to the ER. T-test; ns, not significant.

Selenos is rich in short linear motifs (SLiMs), a common feature of intrinsically disordered proteins (IDPs), which mediate interactions with protein partners^26^. Analysis of selenos’s short linear motifs (SLiMs) by the Eukaryotic Linear Motif (ELM) resource^27^ suggested roles in trafficking, autophagy, and signaling^9^ (Fig. S1). However, the specific proteins that utilize these interaction sites have not yet been identified, despite the extensive number of reported protein partners of selenos^28–30^. A study of selenos’s interactome utilizing affinity purification mass spectrometry showed an abundance of seemingly unrelated multiprotein complexes such as the nuclear pore, oligosaccharyltransferase, coatomer (coats membrane-bound transport vesicles), cyclosome (anaphase-promoting), and ERAD^28^. The large number of interactors and their diverse cellular functions led to the hypothesis that selenos is involved in the maintenance and transport of multiprotein complexes. Yet, this study did not find interactions specific to the redox loop. A separate study of selenos’s interactome using a similar experimental approach identified roles for selenos in metabolism and RNA processing^30^. Both studies emphasize partners whose interactions are strong enough to be captured by affinity purification, whereas proteins interacting with lower affinities are less likely to be detected. Yet these weaker interactors are functionally important for proteins with IDS, such as selenos. Furthermore, while such studies provide a broad overview of a protein’s interaction landscape, they are also unable to differentiate between proteins that interact directly with selenos and proteins that are merely part of a stable complex that also contains the bait protein. Thus, such studies are also not suitable for mapping interacting proteins to their respective SLiM(s) on selenos. Yet, to identify the specific cellular roles of selenos, knowledge of the directly interacting proteins and their interaction site is crucial.

Thus, to identify which proteins are direct protein partners of selenos and determine the partners that utilize selenos’s unusual redox loop, we employ here both global and site-specific crosslinking proteomics, in addition to conventional affinity purification proteomics with different placements of the affinity tag and selenos’s segments (Figs. 1B-D). Comparing and contrasting overlapping data from these three different approaches for recording interactomes, we can pinpoint key selenos partners that directly interact with selenos and thus identify the cellular pathways they likely participate in. Furthermore, we demonstrated that the C-terminal redox loop plays an important role in interactions with proteins involved in translation.

## Results

### Experimental design and rationale

Previous studies of selenos interactomes utilized N-terminal HA^28^ and Flag^30^ tags and C-terminal S-tag^29^. However, the high levels of expressed selenos in these experiments can alter ER morphology and the cytoskeleton^12,32,33^. Thus, we developed stable cell lines allowing us to control expression levels, an essential part of capturing endogenous protein partners by affinity proteomics ^34^. To titrate the level of selenos expression, we used stable expression of selenos under an inducible promoter in Flp-In T-REx 293 cells (a derivative of HEK293). For all experiments, selenos was expressed at levels approximately 60-fold above endogenous levels under non-stress conditions (Fig. S2A), corresponding to about 15-fold above endogenous levels during ER stress (reviewed in^9^). This level of expression did not affect ER or cytoskeletal morphology (Figs. S2C–D).

Another experimental choice was to substitute Sec with Cys at position 188 because the human Sec incorporation machinery has limited capacity and cannot efficiently support overexpression^5^. Such substitutions are commonly employed to improve sample homogeneity by avoiding the subpopulation of truncated selenos resulting from the failed incorporation of Sec *in vivo.* Thus, we used only selenos U188C variants here, but, for brevity, we use selenos and will not explicitly label variants as U188C in every instance. Because of the structural similarity between Cys and Sec^35^, this substitution does not introduce a significant structural perturbation^36^. In endogenous selenos, its Sec forms a selenylsulfide bond with the Cys at position 174, forming an internal loop at the C-terminus. We have previously shown that this redox loop can be “stapled” through a disulfide bond that arises when Sec 188 is substituted by Cys^22,23^. However, since Cys is a poorer nucleophile than Sec, we expect the selenos U188C variant to be a less reactive enzyme. The substitution may also lead to a higher percent of “open” loop because the redox potential of the disulfide bond is higher than that of the selenylsulfide bond.

While the use of small affinity tags is routine in proteomics, both tag placement and sample preparation requires optimization to retain native interactions^34,37^. The nature and placement of the affinity tag warrant especially careful consideration for small proteins such as selenos (189 amino acids). Positioning the tag at selenos’s C-terminus might alter the conformational ensemble of its disordered segment or restrict access to the C-terminal redox loop in which the Sec is positioned. Conversely, a tag at the N-terminus could affect membrane integration or protein partners. The expression and isolation of selenos with different affinity tags were compared, and the V5 tag was selected because it enabled high-yield capture, was compatible with on-bead trypsin digestion, and the digest nanobodies yield fewer peptides than the larger IgGs or proteins used in other affinity resin during trypsin digest. To limit bias and to capture the influence of affinity tag placement, both N-terminally and C-terminally V5-tagged selenos constructs were used in our experiments and compared (Figs. 1B-D). The cellular localization of untagged selenos and selenos with V5 affinity tag was placed on either the N- or C-terminus was imaged using immunofluorescence microscopy, and no discernible differences in their cellular localization were found (Figs. 1F and S2B^10^).

We recorded three types of interactomes: one based on affinity purification, one using global lysine crosslinking, and one using site-specific crosslinking. While each serves its own purpose, a key goal was to combine their complementary information to capture not only strongly interacting partners but also those with weaker, transient interactions. In addition, we aimed to identify the specific selenos segment with which partner proteins interact. Thus, our first workflow was affinity purification sample preparation coupled with data-independent acquisition (DIA) mass spectrometry, which we abbreviate as the non-crosslinked interactome (NC-interactome) (Fig. 1B). Here, we refined previous studies by carefully titrating the levels of stably expressed selenos variants in Flp-In T-REx 293 cells, by using multiple tag positions, and by taking advantage of the increased sensitivity offered by DIA, a sensitive acquisition method. Previous reports of selenos’s interactome used data-dependent acquisition, which captures abundant proteins best. In contrast, DIA identifies more proteins, reduces biases, and enhances reproducibility^38,39^. We recorded the interactomes of selenos with both C and N-terminal V5 tags, and of a selenos variant without the IDS (Figs. 1B and S3A). While the disordered segment spans residues 124-189, we chose to terminate our IDS-less selenos construct after residue 133 to align with a naturally occurring selenos variant that terminates at this position as identified by western blotting (Fig. S3A). This truncation of selenos removed the bulk of the disordered segment, including the segment’s SLiMs as well as the redox loop (Figs. 1A and S1)^33^. Compared to the full-length selenos variants, the truncated V5-selenos 1-133 had an expression level of about 40% and slightly stronger localization to the Golgi (Fig. S2B), which may be due to the removal of a recycling signal that mediates its translocation from the Golgi to the ER. The V5-selenos 1-133 variant was still primarily localized to the ER membrane (Fig. 1F). As a control for our AP-MS-DIA data sets, we employed cells in which an empty expression vector was integrated into Flp-In T-REx 293 cells. Given the complexity of the interactome, five independent biological replicates were utilized to ensure robust statistical analysis.

When optimized, standard affinity purification mass spectrometry isolates strongly bound protein partners and endogenous complexes^34,37^. However, selenos likely participates in both stable and weakly associated complexes^28^, and previous work showed that capturing it together with its partner, the ATPase p97 (low μM affinity ^40^), required crosslinking to prevent selenos from dissociating from the complex during sample preparation^41,42^. Thus, we have employed cross-linking in live cells to supplement the information from the NC-interactomes and expand the coverage of protein partners with lower binding affinity (i.e., higher dissociation constants). To retain such weakly bound partners in our sample and identify proteins that are in the immediate vicinity of selenos, we used global lysine crosslinking in live cells using dithiobis(succinimidyl propionate) (DSP), which can cross the cell membrane to allow crosslinking *in vivo*. Following the isolation of proteins covalently bound to selenos, the DSP linker is reduced to break the covalent association between proteins, allowing improved trypsin access and more efficient proteolytic cleavage by trypsin. At the low concentrations used here (Fig. S3C), it acts as a mild crosslinker and is commonly employed to capture protein assemblies that would otherwise dissociate during sample preparation. It also offers the benefit that a change in enrichment indicates that the proteins was in the proximity of selenos *in vivo*, prior to lysis. Furthermore, global crosslinking is expected to enrich proteins with exposed lysines positioned near exposed lysines on selenos, and therefore the resulting interactome is expected to differ from that of non-crosslinked samples. The distribution of lysines available for crosslinking in selenos is shown in Figure 1E. For comparative mass spectrometry analysis, we employed Flp-In T-REx 293 cells expressing V5-selenos with and without DSP (Figs. 1C and S3C). We chose the N-terminal V5-tagged selenos for global lysine crosslinking because it leaves selenos’s C-termini unmodified and accessible. We refer to this approach as the lysine crosslinking interactome (LC-interactome).

Lastly, identifying the selenos segment responsible for interaction with a given partner provides important biochemical context, revealing whether multiple partners compete for the same site and whether the selenos redox loop is involved. To identify which segments of selenos are responsible for binding specific proteins, and to further trap weakly interacting protein partners and those likely to be direct protein partners (i.e. bound to selenos), we used site-specific insertion of the photocaged crosslinker DiZPK (3-(3-methyl-3H-diazirine-3-yl)-propaminocarbonyl-Nε-L-lysine). This crosslinker resembles lysine^43^ and forms, upon photoactivation, a carbene that reacts with any proteins in close proximity. We call this approach site-specific crosslinking (SC-interactome) (Fig. 1D). For its site-specific insertion into selenos, we chose two positions for different experiments. To probe for interactions with membrane-associated proteins, we chose position A68, which is in the second helix near the membrane interface but still accessible to the cytoplasm. To study interactors with selenos’s redox loop, position R180 was used because it is centrally located within the redox loop (Fig. 1D). The resulting selenos-V5 variants with DiZPK inserted are labeled as A68DiZPK and R180DiZPK, respectively. To compensate for the lower yield resulting from some proteins truncating at the incorporation site of unnatural amino acids, we employed transient expression in HEK293T for these selenos variants. This approach produced levels of DiZPK-incorporated selenos similar to those observed for selenos variants that we stably expressed in Flp-In T-REx 293 cells (Fig. S3D). To identify proteins enriched through crosslinking, samples collected with and without photoactivation were analyzed and compared. The illumination level used for crosslinking was shown not to lead to proteome changes during the illumination period^44^.

The three types of interactomes, NC, LC, and SC, differ from one another. NC will capture strongly bound partners and will have no compositional bias, i.e. it does not depend on the availability of lysines or DiZPK. However, it may allow proteins partners with lower binding affinity (higher kD) such as p97 to exchange during sample preparation. In contrast, LC and SC interactomes reveal proteins enriched through crosslinking in live cells, prior to lysis. Therefore, proteins that bind strongly to selenos but are not enriched (for example, because they are distant from the crosslinking site or lack sufficient lysine residues) will appear at similar levels in samples with and without crosslinking and thus will not be part of the interactome. Consequently, although no single dataset can represent the entire cellular interactome, together they provide the broadest overlap of information.

### Selenos engages with biogenesis of membrane proteins

When comparing the different interactomes, we observed significant enrichment of complexes that coordinate the insertion, folding, processing, and assembly of membrane proteins and their complexes within the ER membrane. Essentially, these complexes not only insert membrane proteins into the lipid bilayer based on hydrophobicity and structural features, but also guide their intramembrane folding and assembly into the appropriate complexes in which they operate^45^. The specific features of the nascent peptide chain determine how the newly synthesized protein is handed off between insertion complexes during translation^45,46^. Insertion and folding of membrane proteins are mediated by several key complexes, including the Sec61 translocon, GEL GET and EMC like), EMC (ER membrane protein complex), PAT (protein associated with the ER translocon), BOS (back of Sec61), and SND (SRP-independent). These complexes are associated with chaperones that could assist in the assembly of complexes as well as factors responsible for sensing aberrant membrane proteins and complexes and directing their recovery or degradation as needed. Glycosylation of membrane proteins can take place co- or post-translationally by the oligosaccharyltransferase (OST), which glycosylates proteins on Asn (N). When relevant, signal peptides can be released from newly synthesized membrane proteins by the signal peptidase complex (SPC)^46^. These complexes physically couple to coordinate membrane insertion, folding, modification, and assembly of membrane proteins in a dynamic fashion, in response to the specifics of the proteins to be inserted and folded^46^. Hence, such ensembles of complexes are not static entities with defined stoichiometry, but rather form a dynamic network of supramolecular assemblies within the ER membrane^47^.

#### · The ER membrane protein complex (EMC)

Among these insertion complexes, components of the EMC are abundant in our interactomes (Fig. 2B), and across all crosslinked interactomes, EMC components are among the most highly enriched proteins (Table S3). The EMC specializes in the insertion of transmembrane domains with low to moderate hydrophilicity^48^, the folding of the proteins post-insertion^49^, and the coordination of complex formation of the inserted proteins with additional components^50,51^. The EMC’s cellular roles extend beyond its contribution to membrane protein insertion, since changes in its levels affect not only protein homeostasis but also processes such as lipid synthesis and organelle communication^52,53^. One confirmed alternate function is the EMC’s involvement in the transfer of phospholipid from the ER to mitochondria at ER-mitochondria contact sites (MAMs)^54^. Although there are 10 EMC subunits, the complex assembles from only 9 at a time, since EMC8 and the less abundant EMC9 are mutually exclusive (Fig. 2B). All 10 EMC components were present in the LC-interactome, possibly because the DSP crosslinked the EMC complex itself. Similarly, eight components were present in SC-interactomes, with EMC7 the most enriched. EMC7 binds EMC2, which is the most enriched EMC component in NC-interactomes. EMC2 is part of the entry path for the client transmembrane domains to be inserted into the bilayer by EMC3^55^. Because EMC2 was detected in all six interactomes, we used it to validate this observation. EMC2-Flagx3 was used to pull down untagged selenos (Fig. S4), and the amount of bound selenos increased following DSP crosslinking.

**Figure 2.**
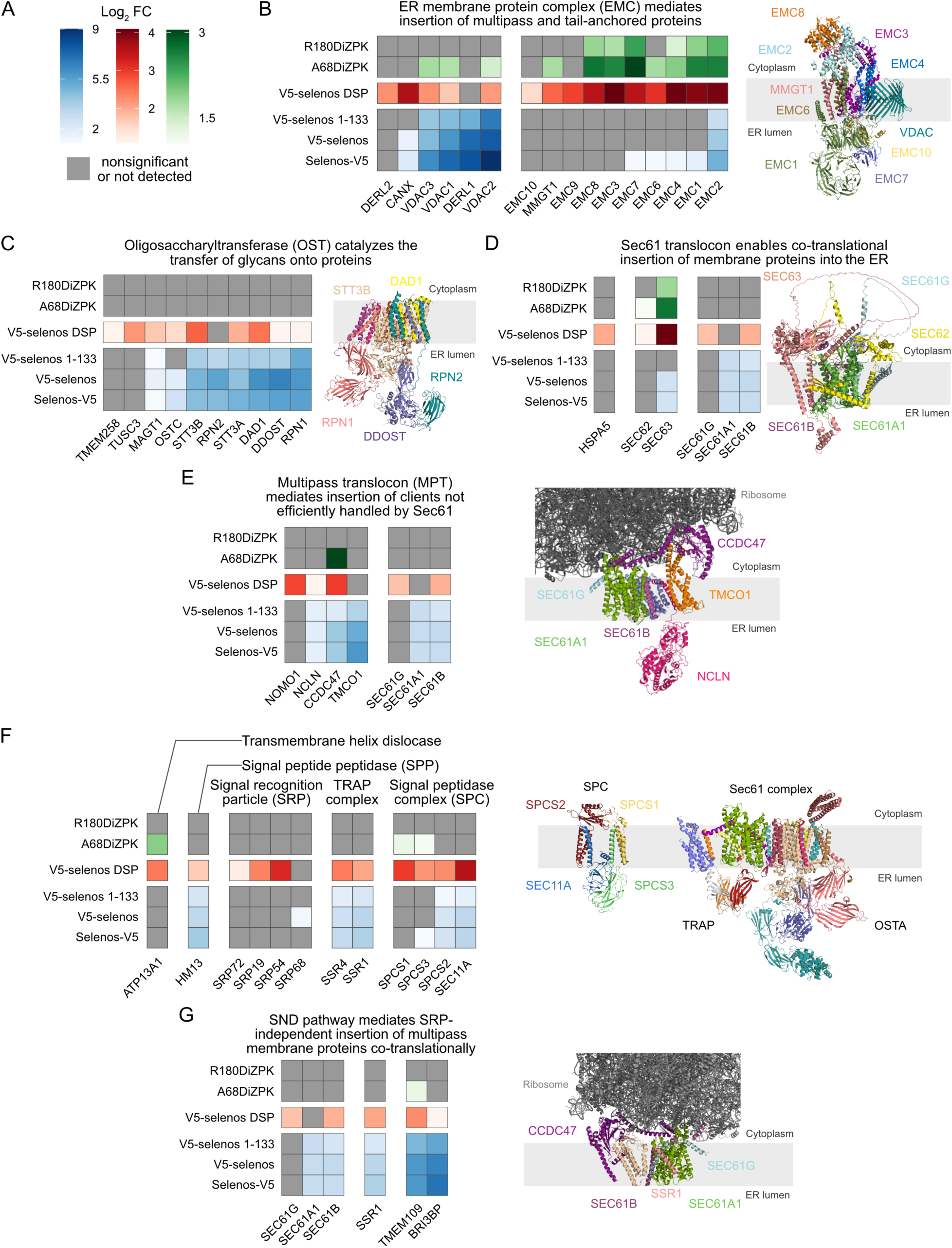
Analysis of proteins in selenos interactomes related to membrane biogenesis. A) The Log_2_-transformed fold change (Log_2_ FC) for all panels: Log_2_ FC from the NC-interactome experiments is shown in blue, from LC-interactomes in red, and SC-interactomes in green. Proteins that are nonsignificant or not detected under a given condition are shown in gray. Significant proteins were defined as those with an adjusted p-value below 0.05 and Log_2_ FC above 1 (Tables S1-3). B) Left: enrichment of significant proteins related to the ER membrane protein complex (EMC) in the selenos interactome. Right: structure of the EMC bound to VDAC (PDB ID: 8J0O). C) Left: enrichment of significant proteins related to the oligosaccharyltransferase (OST) complex. Right: structure of OST-B (PDB ID: 6S7T). D) Left: enrichment of significant proteins related to the Sec61 translocon complex. Right: prediction by AlphaFold3 with several flexible segments removed for clearer visualization. E) Left: significant proteins related to the multipass translocon (MPT) and their enrichment in the selenos interactomes. Right: the structure of these components with a docked translated ribosome (PDB ID: 6W6L). F) Communication between complexes involved in membrane protein biogenesis is shown in the complex of Sec61-TRAP-OSTA translocon complex (PDB ID: 8B6L) and the SPC complex structure (PDB ID: 7P2P). G) Left: enrichment of significant proteins related to the SND pathway. Right: the structure of the ribosome-bound SND3 translocon (PDB ID: 9I78).

Based on the enrichment in the three interactomes, which provide information about the segment of selenos involved, the distribution of lysine residues on the EMC surface, and the known structure of the EMC complex, it is possible to map selenos’s interactions with the EMC complex. (Note that AlphaFold3 models could not be used here because their predictions of multi-protein membrane assemblies often orient selenos incorrectly with respect to the cytoplasm/lumen). It can be inferred that the hydrophobic segment of selenos is located near the EMC3/EMC4/EMC7 subcomplex rather than the MMGT1/EMC6 subcomplex, while the C-terminus of selenos is situated close to the EMC7/EMC2 subunits.

The link between selenos and the EMC, in part, helps explain the high levels of the outer mitochondrial membrane protein VDAC2 (voltage-gated ion channel 2) observed in the selenos interactomes. Cryo-EM studies have shown that EMC purified from human cells contains mitochondrial non-selective voltage-gated ion channels (VDACs)^56^, which transport proteins, ions, and small molecules to the mitochondria^57^. In addition, VDACs also act as lipid scramblases and play a role in autophagy^58^, and, *in vivo*, they interact with the EMC at ER-mitochondria contact sites^56^. When VDACs are present, EMC3 undergoes a conformational change that occludes its hydrophobic vestibule, thereby preventing newly synthesized proteins from exiting EMC3 to the membrane. This led the authors to suggest that contact between VDACs and the EMC may be related to functions other than protein insertion, such as lipid exchange between the ER and mitochondria^54^. When EMCs are not bound to VDACs, they distribute throughout the ER membrane^56^. It is known that VDACs also interface, like selenos, with the EMC3/EMC4/EMC7 subcomplex (PDB 8J0O^56^, Fig. 2B). This places them near to each other and possibly close enough to enrich VDACs in the SC-interactome of selenos (Fig. 2B and section about mitochondrial proteins). VDACs are further discussed in a subsequent section.

Also at high enrichment are the EMC partner derlin-1^29,59,60^ (Fig. 3B), which is a well-established interacting partner of selenos^29,59,61^. Derlins belong to the rhomboid intramembrane protein family^62,63^, whose members specialize in the recognition of membrane proteins inside the lipid bilayer. Specifically, derlins facilitate the retrotranslocation of misfolded membrane proteins from the ER into the cytosol for proteasomal degradation^64,65^. The role of derlins in relation to the EMC is unknown, but based on their yeast homolog, they may be used to hold orphan protein partners as a chaperone until other partners are available or until stress is resolved^66^. They can also serve as a quality control factor as part of their ERAD-related roles, as will be later discussed.

**Figure 3.**
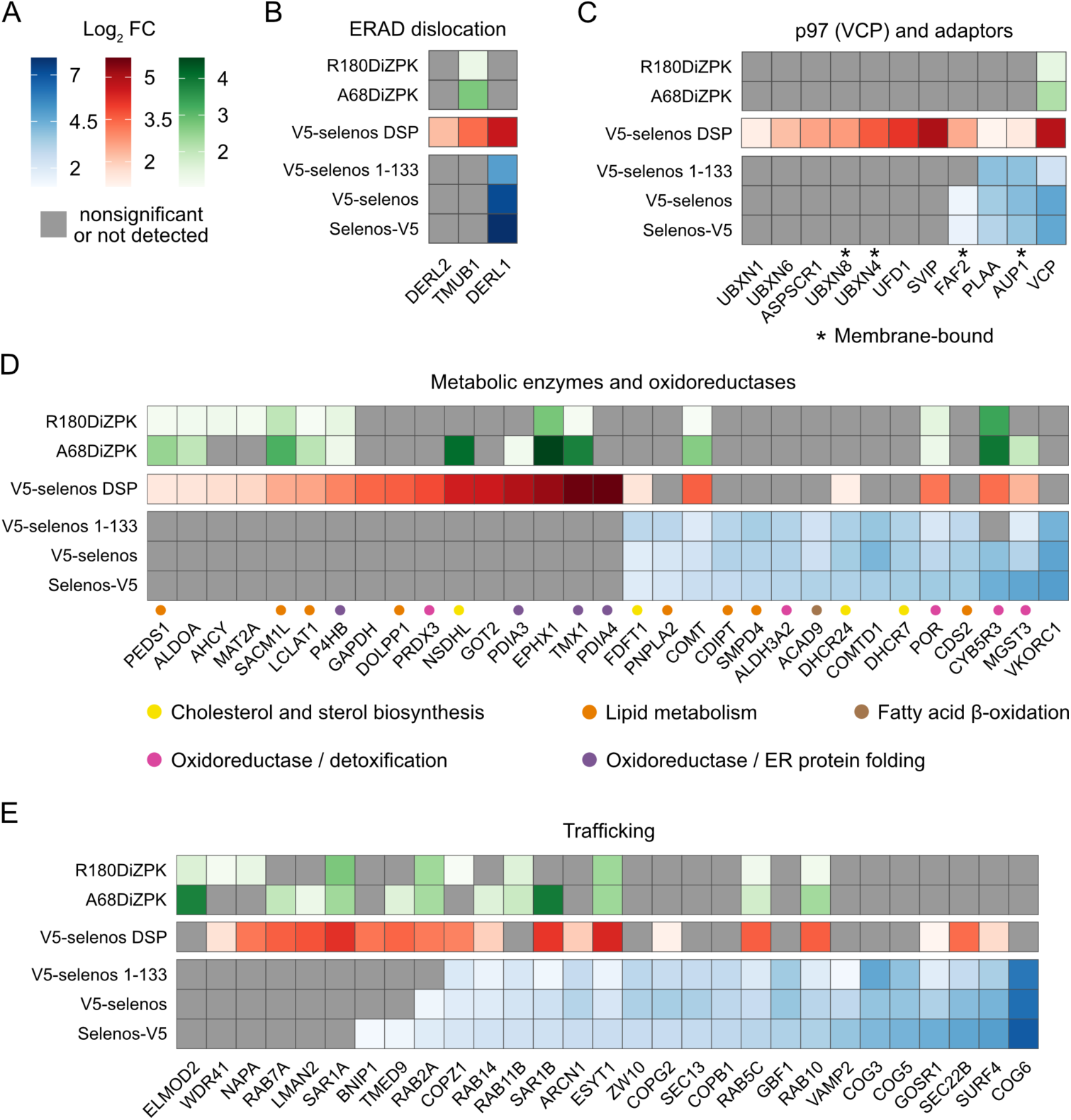
Selenos interactome in the context of derlins, p97, and metabolism and protein trafficking. A) The Log_2_-transformed fold change (Log_2_ FC) for all panels: Log_2_ FC from the NC-interactome experiments is shown in blue, from LC-interactomes in red, and from SC-interactome in green. Proteins that are nonsignificant or not detected under a given condition are shown in gray. Significant proteins were defined as those with an adjusted p-value below 0.05 and Log_2_ FC above 1 (Tables S1-3). B) ERAD dislocation factors. C) p97 and its adapters. D) The top ten highly enriched metabolic enzymes and oxidoreductases in NC- and XL interactomes are shown. E) The top ten highly enriched trafficking associated proteins in NC- and XL interactomes are shown.

#### · The oligosaccharyltransferase (OST)

The oligosaccharyltransferase (OST) is another membrane-embedded complex within the membrane protein biogenesis supramolecular assembly, several components of which are highly enriched across the different selenos interactomes. Its enzymatic activity is the transfer of glycans to proteins^67,68^ (Fig. 2C). The OST complex is anchored in the ER membrane, with most of its volume extending into the ER lumen where its catalytic site resides. The complex exists with two distinct compositions. OST-A contains the catalytic dolichyl-diphosphooligosaccharide protein glycosyltransferase subunit STT3A and inserts glycans co-translationally. OST-B, on the other hand, contains the catalytic dolichyl-diphosphooligosaccharide protein glycosyltransferase subunit STT3B, which does not interact with the ribosome. OST-A glycosylates nascent polypeptide chains during translation, while OST-B glycosylates proteins post-translationally^68,69^. Components of OST-A and OST-B were already noted in studies of selenos’s interactome^28^. However, the interactomes here allow us to identify the most likely location of selenos in the complex.

NC-interactomes show the highest enrichment of the OST components DDOST (dolichyl-diphosphooligosaccharide-protein glycosyltransferase 48 kDa subunit, also known as OST48), RPN1 (dolichyl-diphosphooligosaccharide-protein glycosyltransferase subunit 1), and DAD1 (dolichyl-diphosphooligosaccharide-protein glycosyltransferase subunit). There is no enrichment in SC-interactome as one would expect because the OST complex is primarily embedded in the ER membrane and lumen, while A68 and R180 are cytoplasmic. The resulting lack of site-specific interactions between OST components and selenos reflects, therefore, the structural organization of the OST complex rather than the degree of interactions. In the NC-interactome STT3B and DAD1 are most enriched; of course, this reflects not only proximity to selenos but also the lysine distributions on the surface of selenos and partner proteins. Even though the N-terminus of selenos can react with DSP, it is more likely that selenos’s Lys50, which lies in close proximity to the cytoplasmic membrane interface, is responsible for lysine crosslinking with OST components, given the presence of a cluster of lysine residues in STT3A/B and DDA1 at this same interface. DDOST is bound to DAD1 (Fig. 2C), which explains its enrichment in NC-interactomes.

There is no reported evidence that selenos influences glycosylation by OST. Nevertheless, we tested whether glycosylation of α1-antitrypsin (A1AT) aberrant New Hong Kong (NHK) variant, which is a model ERAD substrate, is affected by the absence of selenos (Fig. S5). No significant difference was observed. Indeed, while selenos is tightly anchored to OSTA/B, most of its mass is cytoplasmic and is removed from the OST catalytic core. It is more likely to act as a structural protein or a scaffolding protein bridging OST with other complexes. When considering the underlying reason for the tight association of selenos with OST, it is notable that a recent study reported that OST components associate with ERAD substrates and that OST influences ERAD in processes that require derlin-1 but do not involve glycosylation^70^.

#### · The Sec61 translocon and its interacting complexes

The Sec61 translocon is the protein-conducting channel of the insertion machinery, that allows membrane proteins to fold and be released into the lipid bilayer^71^. Functionally, the three subunits form Sec61αβγ, the membrane channel, while Sec62 controls gating from the channel to the membrane^72^, and Sec63 governs the docking of the ribosome^73^. Components of the Sec61 translocon, the protein-conducting channel of the insertion machinery, are enriched in the selenos interactomes. The three subunits of the Sec61 translocon, Sec61αβγ, are present at moderate levels with and without crosslinking (Fig. 2D). Sec63 stands out with high levels in the LC and SC crosslinking interactomes. This suggests that selenos cytosolic helix passes near Sec63 or they would not be able to be crosslinked in a site-specific manner. It likely interacts with selenos’s IDS since it is absent in V5-selenos 1-133 NC-interactome, but it is trapped efficiently by both A68DiZPK and R180DiZPK. Also present is the chaperone HSPA5 (BiP), which interacts with Sec62’s luminal segment and drives unidirectional translocation by ATP-dependent binding and release cycles.

The core Sec61αβγ channel associates with several complexes beyond Sec62/63^74,75^. Furthermore, there is evidence that complexes associate with the Sec61 translocon to form a lipid filled cavity behind Sec61 in which proteins can fold without directly relying on Sec61^76^. To simplify the discussion of components observed in this study, their function and subunits are listed in Fig. S6. Of those, OST and EMC were already discussed above. Additional complexes whose components are significantly enriched in both NC, LC, and SC are the GEL and PAT. Especially, TMCO1 (calcium load-activated calcium channel) which is part of the GEL, and then CCDC47 (PAT complex subunit CCDC47). These PAT and GEL components are bound to the Sec61αβγ^72^ (Fig. 2E). The most enriched proteins are TMCO1 (calcium load-activated calcium channel), which is part of the GEL, and then CCDC47 (PAT complex subunit CCDC47). These PAT and GEL components are bound to the Sec61αβγ^72^ (Fig. 2E). TMCO1 was identified as a member of the Oxa1 family, which is responsible for the insertion of membrane proteins^77^ and may have the ability to insert membrane proteins independently when assembled with the Sec translocon. EMC3 is also a member of the family. CCDC47 is a protein partner of the transmembrane asterix and the transmembrane TMCO1. (Asterix, which is an obligate partner of CCDC47, is typically not detected in the proteomics studies because it is resistant to trypsin digestion and yields few peptides^73,76^, and it was therefore not observed in the selenos interactome.) Both TMCO1 and CCDC47 directly interact with docked ribosomes (PDB 6W6L). However, those complexes may not be involved in membrane protein biogenesis and insertion of membrane proteins. CCDC47 was previously shown to associate with derlin-1, derlin-2, and selenos in both HEK293 and mouse enterocyte cells^78^. Its knockdown in HEK293 reduced the degradation of the model ERAD substrate null Hong Kong α1-antitrypsin (NHK-A1AT). It is intriguing that once again we find that the components most enriched in crosslinking are those that bind the ribosome (see later).

Additional complexes known to interact with the Sec61 translocon and whose components are present at lower enrichment levels (Fig. 2F) are: the translocon-associated protein (TRAP)^79^, the signal recognition particle (SRP) complex, signal peptidase complex (SPC), and the signal peptide peptidase (SPP). SRP recognizes the N-terminal signaling peptide of precursors of secreted or transmembrane proteins and directs the nascent chain of the transcribed protein on the ribosome. Incidentally, it is known that derlin-1 associates with the SRP only when the unfolded protein response (UPR) is activated^80^. SPC and SPP remove signal peptides (Fig. 2F). SPP was shown to interact with derlin-1^81^. Also present is the ATP13A1 endoplasmic reticulum transmembrane helix translocase, a P-type ATPase that removes misdirected mitochondrial membrane proteins from the ER membrane^82^. It engages with the Sec61 translocon for handling substrates^83^. Since enrichment of these additional complexes is rather low in NC-interactomes and is most prominent in the LC-interactome, they are less likely to be direct partners. Instead, crosslinking of the supramolecular complexes near the Sec61 translocon enables their retention in LC-samples.

Not all the complexes discussed can be bound to Sec61αβγ simultaneously, and it was shown, for example, that PAT excludes the binding of the OST^72^. Hence, selenos emerges as interacting with several complexes that communicate with the Sec61 translocon.

#### · The SRP-independent (SND) pathway

The SND pathway is a parallel pathway to the SRP pathway. SND3, in humans TMEM109 (voltage-gated monoatomic cation channel TMEM109; also called mitsugumin 23)^84^, acts as the insertase. A recent structure of the SND complex from *Chaetomium thermophilum*^85^ showed coordination between Sec61αβγ, TRAPα, and CCDC47 (Fig. 2G). TMEM109 exhibits high enrichment levels in all NC-interactomes and is also found in LC and SC interactomes. Also present is the TMEM109 paralog, BRI3BP (BRI3-binding protein). Information about BRI3BP is sparse^86^. In contrast, TMEM208 (transmembrane protein 208, Snd2), which is considered TMEM109 partner^87^ is absent.

In summary, the various selenos interactomes and respective protein enrichments indicate that multiple complexes involved in the biogenesis, processing, modification, and quality control of membrane proteins are localized in the vicinity of selenos. The crosslinking patterns support the notion that selenos directly interacts with the OST, EMC, and components of the multipass Sec61 translocon, such as PAT, SND, and Sec63. Notably, the most enriched components in all these different complexes constitute the insertase and proteins that coordinate ribosome engagement.

### Selenos known partners, derlins and p97, are relevant for specific complexes

While selenos has been extensively investigated in the context of the ERAD pathway^88–91^, conflicting evidence has left its precise role unresolved. However, there is consensus that of the ERAD components, the soluble AAA+ ATPase p97 (VCP)^14,40,92^ and the transmembrane derlins^93^ are consistently found to bind and collaborate with selenos.

The derlins, which were introduced above, have in addition to their established roles in protein quality control, roles in cholesterol biosynthesis^94,95^, intracellular signaling pathways^96^, and the regulation of autophagy^97^. While the three human derlin paralogs, derlin-1 (DERL1), derlin-2 (DERL2), and the less abundant derlin-3 (DERL3) share sequence homology, they are members of different protein complexes and execute non-redundant functions^95^. In our NC-interactome experiments, derlin-1 is among the most enriched proteins, while derlin-2 is below the threshold of significance (Fig. 3B, Table S1). In our LC-interactome experiments, both derlin-1 and derlin-2 were enriched. However, despite containing fewer lysines, derlin-1 showed a higher level of enrichment. Overall, derlin-1 is corroborated as a top selenos interactor in unstressed cells. However, this is not the case for derlin-2.

Derlins are classified as ERAD dislocation factors because they assist in the dislocation of proteins with transmembrane helices into p97 on the path to degradation. In addition to derlins, the dislocation factor, TMUB1 (transmembrane and ubiquitin-like domain-containing protein 1), a membrane protein that, similar to derlins, recognizes unfolded clients and delivers them to p97^98^. TMUB1 is present only in LC- and SC-interactomes (Fig. 3B).

The association between selenos and derlin is attributed in the literature to their collaboration in ERAD complexes^93^. Our interactome analysis identified, in addition to derlins and p97, several components of the ERAD machinery, but their number and enrichment levels are low (Fig. S7A and B). This low enrichment suggests that they interact indirectly with selenos (i.e. through other protein partners or complexes). Our experiments were carried out without stressors or unfolded protein clients, so it is possible that selenos is recruited to ERAD complexes upon stress or when clients are abundant. Nevertheless, without stress, these results do support the presence of selenos at high levels in multiple ERAD complexes. The supporting information contains additional figures displaying ERAD partners present in the interactome at low levels (Figs. S7A and B) and the composition of ERAD complexes^93^ that were identified in the literature as containing selenos.

In contrast, there are multiple enzymes that are linked to cholesterol and lipid biosynthesis, an emerging cellular role of derlins^95^. Figure 3D demonstrates that the enzymes found in selenos interactomes can be classified as redox enzymes, cholesterol/sterol biosynthesis, lipid biosynthesis, and oxidoreductases. We note that in Fig. 3D, many enzymes that have crosslinked in the A68DiZPK SC-interactome are also crosslinked to R180DiZPK in the SC-interactome. The link between selenos and metabolism is corroborated by a rich literature about selenos in metabolic disorders^9^. A study of a hepatocyte-specific SELENOS knockout mice^17^ demonstrated that total cholesterol is higher in the KO in high-fat diet (HFD). Therefore, at least in relation to cholesterol in serum, selenos acts as a brake on cholesterol synthesis or secretion. Additionally, selenos knockout influences the level of several metabolic enzymes. Among these enzymes is fatty acid binding protein 1 (FABP1), which contributes to metabolic disorders, and whose levels were shown to be regulated by derlin-1 linked degradation^99^.

Also found at significantly higher enrichment levels than ERAD components are trafficking proteins (Fig. 3E). Most trafficking-related components are related to ER to Golgi and Golgi to ER trafficking. It is however, hard to assess the link between selenos and trafficking because it was already shown that selenos’s influence on trafficking may be linked to the ER shape and microtubules-dependent processes^33^. In our study, there is a notable level of small GTPases and their GTPase-activating proteins in the SC-interactomes. Finally, we note that while the relationship of selenos to trafficking is unclear, it may involve derlin, which relies on SURF4 (Surfeit locus protein 4) for some of its degradation substrates^100^. Lastly, selenos contains SLiMs related to trafficking (Fig. S1) and hence is able to recruit relevant proteins.

Due to the overlap between derlin-1’s role in membrane protein biosynthesis, metabolism, ERAD, and trafficking, there is a significant overlap between the different selenos interactomes reported here and a larger cluster of ER-related proteins, including selenos and derlin, which was isolated when protein complexes were characterized using size exclusion and identified by subsequent proteomics (Fig. S8)^101^. Overall, distinguishing proteins by assigning them to a single functional category (e.g., metabolism, trafficking, or membrane protein biogenesis) is somewhat arbitrary. It may be more accurate to describe selenos interactions with derlins as central to their function within this ER protein hub. Moreover, based on interactomes obtained under non-stress conditions, the association with derlins appears to extend beyond ERAD and contribute to roles in lipid biosynthesis and cholesterol homeostasis.

### Selenos binds the unfoldase p97 in specific complexes

Selenos and derlins bind p97^12,13,32,102,103^, an ATPase responsible for disentangling and unfolding misbehaved proteins^104,105^. In addition to selenos and derlins, p97 utilizes at least 30 other adaptors that determine its recruitment to distinct cellular locations and processes, and define which client proteins will be identified and processed^106^. Selenos/p97 interactions were previously reported in proteomics studies of p97 adaptors, although selenos’s cellular level is considerably lower than that of other adapters^41,42,107^. Here, p97 is observed as a selenos binding partner under all conditions tested (Fig. 3C). p97 enrichment in V5-selenos 1-133 NC-interactome is lower than that for the full-length variant because selenos’s C-terminal loop contributes to p97 binding^14^. The level of p97 in the NC-interactome may be underestimated relative to its extent of interactions *in vivo*, because only ATP-bound p97 binds selenos^13^, and ATP bound to p97 is likely to hydrolyze during sample preparation, leading to dissociation from selenos. In contrast, the LC-interactome would capture their *in vivo* interactions since both p97 and selenos are rich in lysines.

Determining which p97 adaptors are present across the selenos interactomes obtained using the different approaches should help clarify two important questions: Firstly, after selenos binds, can p97 still recruit any additional adaptors? Secondly, given the p97 adaptors contained in the selenos interactomes, which protein complex(es) of the ERAD is selenos residing in? In the NC-interactome, only three p97 adaptors are appreciably enriched (Fig. 3C). The first, AUP1 (ancient ubiquitous protein 1, also called lipid droplet-regulating VLDL assembly factor) is a membrane-bound p97 adapter. It facilitates the degradation of proteins relevant for the regulation of lipid droplets^108^. It is part of an HRD1-centered complex that associates with derlins, but only a few other components are detected in either the NC- or crosslinked interactomes. AUP1 also contributes to innate immunity^109^ in processes that involve derlin-1 (Fig. 3B) and signal peptide peptidase (Fig. 2F). The second p97 adapter is the phospholipase A-2-activating protein (PLAA or PLAP), which binds the C-terminus of p97, the exit pore from which unfolded p97 clients emerge^110^. Its cellular roles includes regulation of P-bodies^111^, vesicle recycling^42,112^, and the degradation of mitochondrial proteins by the AAA ATPase ATAD1^113^. The third p97 adaptor is FAF2, known as ubiquitin regulatory X domain-containing protein 8 (UBXD8). It is an ER membrane-associated protein that also has roles in lipid droplets^114^ and the mitochondria-ER contact sites^115^. In addition, FAF2 regulates fatty acid synthesis and triglyceride metabolism^114,116^, as well as autophagy-based degradation of peroxisomes (pexophagy)^116–118^ or mitochondria (mitophagy)^115^. FAF2 was previously shown to interact with selenos^29^ and is also present in both derlin-1 and derlin-2 interactomes based on global mapping of cellular protein complexes^59^.

Association of adapters with p97 is highly dynamic and conventional immunoprecipitation can reflect the exchange during sample preparation^41^. In contrast, the LC-interactome traps physiological p97 complexes *in vivo* and prevents exchange during purification. Specifically, LC-interactome enables the capture of proteins that bind directly to selenos, as well as proteins bound to p97 when p97 is associated with selenos. However, even in the LC-interactome, the number of p97 adaptors other than selenos is still low (Fig. 3C). Among the additional adaptors that appear solely in the LC-interactome is the small VCP/p97-interacting protein (SVIP)^119^, which is necessary for lysosomal homeostasis^120^, as it recruits p97 to lysosomes, ensures their dynamic stability, and autophagosome-lysosome fusion. This implies that p97 can be bound by both selenos and the SVIP. Also present in the LC-interactome is the highly abundant UFD1 (ubiquitin recognition factor in ER-associated degradation protein 1), which is one of the most abundant p97 adaptors in the cell and cooperates with NPLOC4 (NPL4; nuclear protein localization protein 4 homolog; not present in selenos interactome) to initiate unfolding of the initiator ubiquitin by p97. Together, they direct the polyubiquitin chain into the central pore of p97 for processing^121^. In addition, there is a UFD1 microvariant version that plays a role in stress response^122^. UBXN4 (UBXD2; UBX domain-containing protein 4) is the other significantly enriched adaptor of p97, which is also an adaptor of the mitochondrial AAA+ ATPase ATAD1^113,123^. ATAD1, which is present in the selenos interactome in both NC- and LC-interactomes (Fig. S7D), is involved in the degradation of membrane proteins^124^.

Consequently, we note the absence of some of the most abundant p97 adaptors especially those in ERAD processes. It appears that selenos collaborates with p97 in specific complexes that include a subset of other adaptors.

### A VDAC-related function for selenos is suggested by the presence of mitochondrial proteins in its interactomes

Additional observation is the enrichment of mitochondrial proteins present in all three types of interactomes (Fig. 4B). The three outer membrane voltage-dependent channels VDAC2, VDAC1, and VDAC3 are present in all three interactomes, with VDAC-2 most enriched. Notably, in our interactomes, some of the most enriched mitochondrial components bind VDACs. A case in question is CYB5R3 (NADH-cytochrome b5 reductase 3), which is highly enriched in crosslinked interactomes. CYB5R3, which is localized to both the ER and the mitochondrial outer membrane (MOM), is a flavoprotein that transfers electrons from NADH to FAD to substrates^125,126^. It supports VDACs functionality by maintaining a locally reduced environment near VDACs^126^. CYB5R3 is in a hub of proteins whose levels are influenced by silencing of selenos in CD4+ T cells^19^. We note that because it is missing in the V5-selenos 1-133 interactome and highly enriched in the R180DiZPK SC-interactome, interactions with selenos must require the IDS.

**Figure 4.**
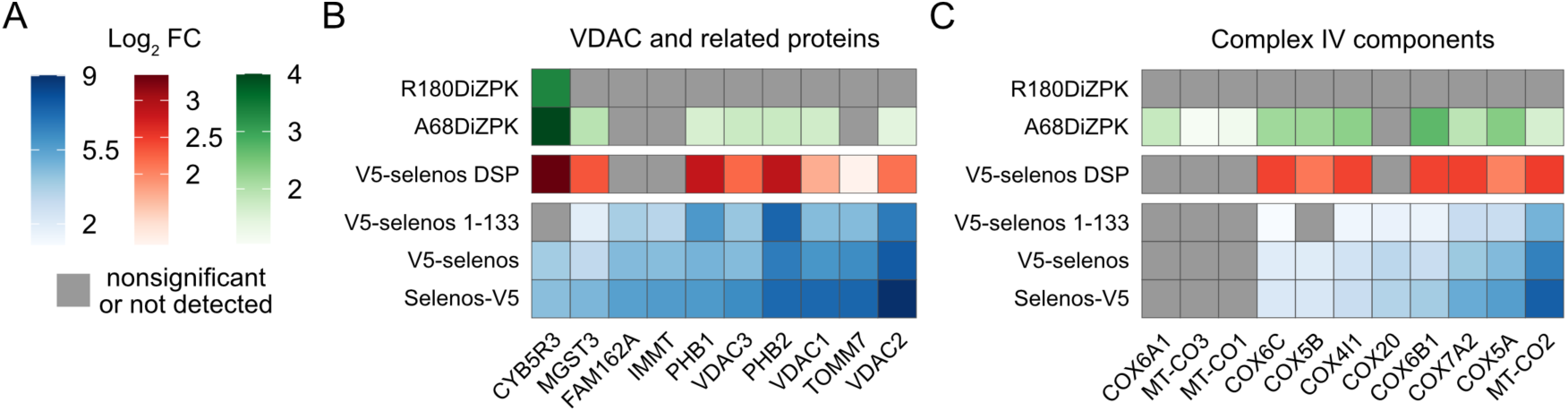
Mitochondrial proteins in the selenos interactome. A) The Log_2_-transformed fold change (Log_2_ FC) for all panels: Log_2_ FC from the NC-interactome experiments is shown in blue, from LC-interactomes in red, and from SC-interactome in green. Proteins that are nonsignificant or not detected under a given condition are shown in gray. Significant proteins were defined as those with an adjusted p-value below 0.05 and Log_2_ FC above 1 (Tables S1-3). B) Highly enriched mitochondrial proteins, selected using the MitoCarta 3.0 database^135^. The list predominantly contains membrane proteins, many of which associate with VDACs. Several proteins are known to be present in additional compartments, including the ER membrane. C) Components of the inner membrane respiratory complex IV (cytochrome c oxidase).

Other proteins in the interactomes are related to VDACs. The prohibitins (PHB1 and PHB2) form a large complex in the mitochondrial inner membrane^127,128^ that cooperates with VDACs. Among the cellular roles of prohibitins^129^ is the degradation of membrane proteins by the membrane bound mitochondrial proteases^130^ and protein quality control^131^. Curiously, it is only PHB2 that is highly enriched in both NC- and LC-interactomes. TOM7 (TOMM7), part of the translocase of the outer membrane (TOM), is enriched in the NC-interactome. A recent cryoEM structure showed that TOM7 is directly bound to VDACs^132^. Furthermore, in yeast, it was shown that the EMC forms a tether between the ER and the mitochondria by associating with the TOM complex. The connection of VDACs to the EMC was already detailed and discussed in relation to Fig. 2. IMMT (MIC60) is part of the MICOS complex, which forms cristae junctions^133^. MICOS is in contact with both VDACs and TOM. MICOS itself associates with FAM162A, also in the NC-interactome, which functions in mitochondrial apoptosis. Lastly, we note among enriched mitochondrial proteins the presence of yet another redox enzyme like CYB5R3 that binds VDACs and influence their activity, MGST3 (glutathione S-transferase 3). MGST3 regulates eicosanoid and glutathione metabolism and its knockdown in cells inhibited apoptosis^134^.

Additionally, multiple components of the inner membrane respiratory complex IV (cytochrome c oxidase) are in the LC- and SC-interactomes (Fig. 4C). Hence, selenos was in close vicinity to these complexes *in vivo*. To a lesser extent and primarily in NC-interactomes we find ATP synthase components and complex I (NADH: ubiquinone oxidoreductase), as well as their assembly factors (Fig. S9D). Affiliated complexes such as ATP synthase are present at low enrichment and are likely indirect interactors, extracted through associations with other mitochondrial inner membrane complexes (Fig. S9B). The PHB1/PHB2 complex also coordinate these complexes^131^.

The enrichment of mitochondrial components in interactomes captured by crosslinking in live cells confirms that these interactions occur within their native biological context. Thus, it seems that a fraction of selenos localizes to mitochondria. Selenos is not known to reside in mitochondria according to MitoCarta 3.0^135^. However, because selenos is present at low levels in the cell (even lower if selenium is not supplemented in the growth medium^136^), if only a small fraction of selenos localizes to the mitochondria relative to its total cellular distribution, its detection would be challenging. It was reported that when selenos is deleted, calcium flux, regulation of reactive oxygen species, and mitochondrial function are impacted^137^.

Because multiple proteins in selenos interactome are present in mitochondria-ER contact sites (MAM)^138,139^, we examined whether selenos may be a tether between the ER and mitochondria. Proteins that act as tethers can increase contact area upon overexpression^140^. Hence, we tested if selenos may be a tether by measuring the contact area between ER and mitochondria in selenos KO HEK293 cells relative to parental cells but found no statistically meaningful change (Fig. S10).

### Selenos’s redox loop interacts with proteins involved in translation

When Gene Ontology (GO) analysis for biological processes was performed using Metascape^141^, for the three NC-interactomes, translation- and ribosome-related terms were enriched for the V5-selenos NC-interactome but not for the selenos-V5 or V5-selenos-133 NC-interactomes (Figs. 5A and S11). In contrast, other biological processes showed similar p-values for all three NC-interactomes. Examination of previously published selenos interactomes with N-terminal HA and Flag affinity tags^28,30^ recorded using different isolation workflows and DDA detection, revealed that they also contain similar biological processes as those found here, predominantly for the V5-selenos NC-interactome data set. Here, however, these processes are substantially reduced in both the selenos-V5 and V5-selenos 1-133 NC-interactome datasets. This implies that positioning the affinity tag at either the N- or C-terminus, as well as the presence of the IDS, may provide insight into the biological roles of selenos’s redox loop (residues 174-188). This observation motivated us to examine in greater detail whether this trend reflects a genuine biological difference due to the presence of the IDS and the accessibility of the redox loop.

**Figure 5.**
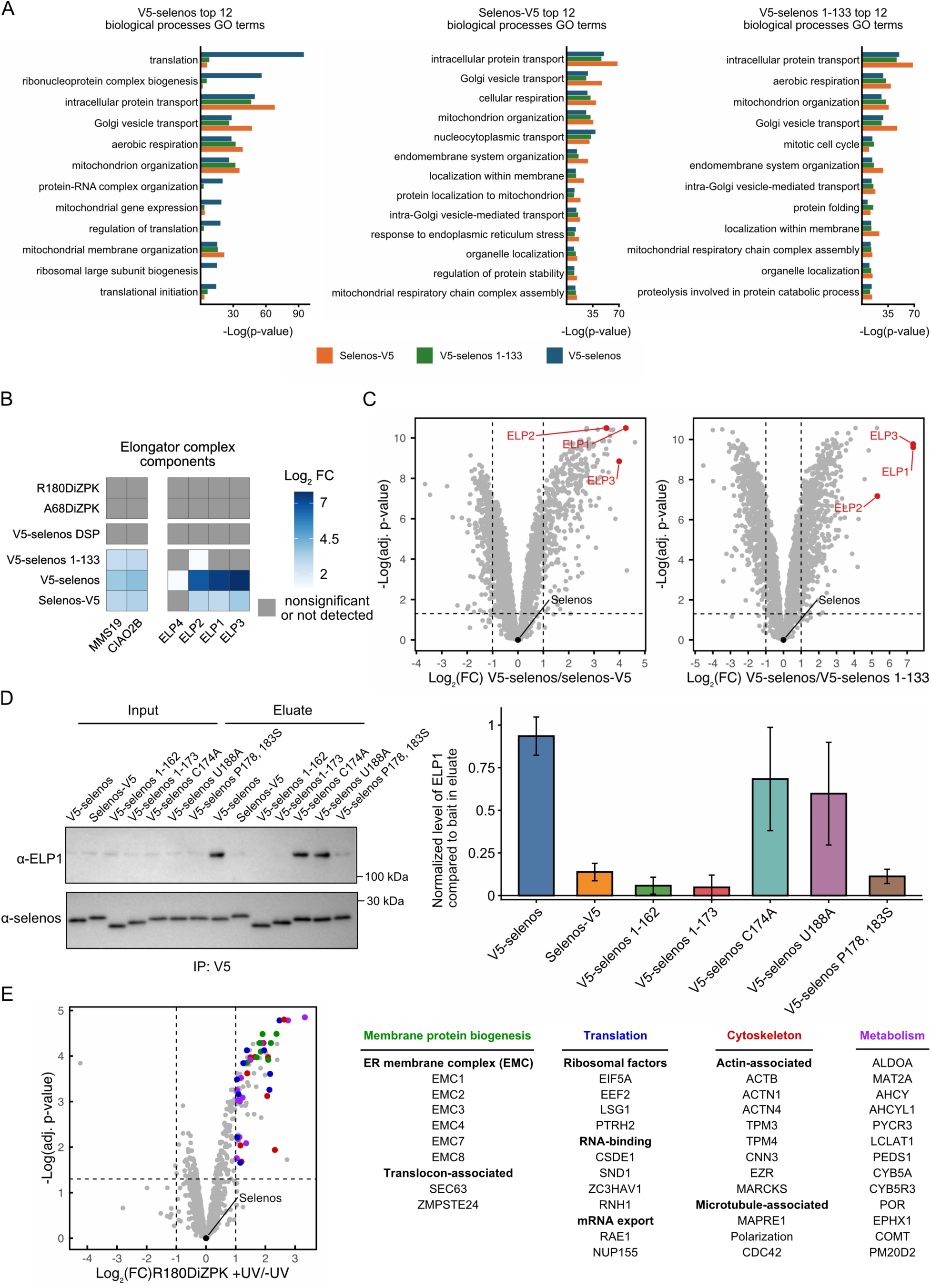
The interactome of selenos is rich in proteins involved in translation. A) Gene Ontology (GO) enrichment analysis of biological processes by Metascape^141^ for NC-interactomes. The top 12 groups (based on p-value) from the NC-interactomes were compared with each other (for the full analysis see Fig. S11). B) Heatmap showing all observed elongator complex components and associated proteins The Log_2_-transformed fold change (Log_2_ FC) for all panels: Log_2_ FC from the NC-interactome experiments is shown in blue. Proteins that are nonsignificant or not detected under a given condition are shown in gray. Significant proteins were defined as those with an adjusted p-value below 0.05 and Log_2_ FC above 1 (Table S1). C) Volcano plots of ELP1, ELP2, and ELP3 in the V5-selenos NC-interactome compared to either the V5-selenos 1-133 or the selenos-V5 interactomes show a strong enrichment in V5-selenos (Tables S4 and S5). D) Variants of the selenos redox loop used to assess their contribution to protein partner binding. Unless otherwise noted, the variants carry a U188C change. ELP levels in eluates were measured using Fiji from three biological replicates. The bar graph shows the mean normalized intensity for each group, with error bars representing the standard deviation. For the scheme introducing the variants, refer to Fig. S16B. E) Volcano plot of proteins enriched in the crosslinked condition for the R180DiZPK variant compared to the non-crosslinked control condition (Table S3). Proteins associated with membrane protein biogenesis, translation, the cytoskeleton, and metabolism are highlighted in the plot.

#### · The elongator complex

Prominent among the proteins involved in translation and among the most enriched proteins in the NC-interactomes (Fig. 5B) are components of the elongator complex subcomplex, ELP1/ELP2/ELP3. The elongator complex is responsible for modifications of the uridines in the tRNA anticodon wobble position^142^. Such modifications have a profound impact on translation fidelity and are critical to elongation regulation^143^. ELP1/ELP2/ELP3 were also present in Turanov’s interactome study^28^, and ELP2 was noted as an upregulated gene when selenos was knocked down in CD4+ effector T cells^19^. When selenos’s IDS is missing (V5-selenos 1-133) or the affinity tag is on the C-terminus of selenos (selenos-V5) there is a significant reduction in ELP1/ELP2/ELP3 (Fig. 5C). We have corroborated that high levels of EPL1 are present in pull-downs with V5-selenos compared to selenos-V5 or variants lacking the IDS, using affinity purifications via V5 nanobodies and western blotting (Fig. 5D). When comparing the levels of ELP1 and selenos in eluates, binding to selenos-V5 is lower, and there is little binding to selenos variants that lack the redox loop. Similarly, variants with an “open” loop conformation, such as V5-selenos C174A or V5-selenos U188A, interact similarly to V5-selenos. However, when the redox loop conformation was altered by changing its two prolines to serines (V5-selenos P178,183S), the variant was also unable to bind ELP1 as efficiently as V5-selenos or V5-selenos variants with an “open” loop conformation. We also confirmed this observation using an HA affinity tag, demonstrating that it is independent of tag identity (Fig. S12). Thus, ELP1/ELP2/ELP3 require selenos’s redox loop for effective binding to selenos (Fig. 5D).

Also, in our NC-interactome, two accessory proteins that assist the elongator are present: MMS19 (nucleotide excision repair protein homolog) and CIA2B (cytosolic iron-sulfur assembly component 2B), which hand off the iron-sulfur cluster to the catalytic ELP3^144^. Their enrichment level in the selenos-V5 and V5-selenos 1-133 NC-interactomes was comparable, indicating that selenos residues 1-133 are involved in their recruitment. Yet their level in the V5-selenos NC-interactome remains higher, suggesting that the IDS still plays a role in binding (Fig. 5B).

#### · Translation initiation factors

Several eukaryotic translation initiation proteins and related regulators are also selectively enriched in the V5-selenos NC-interactome compared with the selenos-V5 and V5-selenos 1-133 NC-interactomes (Fig. S13). There are several components of the translation initiation factor 2B (eIF2B) complex that facilitate the nucleotide exchange of the general translation initiation factor eIF2 to initiate translation^145^. eIF2B does not bind ribosomes directly, and its activity is tightly controlled by the integrated cellular response. Components of the eukaryotic translation initiation factor 3 complex are also present in the NC-interactomes, but at lower levels than the components of the eIF2B complex. In both the LC- and SC-interactomes, there is high enrichment of eukaryotic translation initiation factor 5A-1 (eIF5A). eIF5A coordinates the positioning of the aminoacyl-tRNA on the ribosome to achieve optimal alignment for peptide bond formation^146^. It also promotes termination in stalling ribosomes and regulates the UPR. The high level in the R180DiZPK SC-interactome shows that eIF5A interacts with selenos’s redox loop (Figs. 5E and S13). These interactions demonstrate a link to translation and again suggest that the IDS is involved.

#### · Ribosomes

Additionally, we find high levels of ribosomal proteins in one but not the other two NC-interactomes. Complexes involved in membrane protein biogenesis (see discussion above) interact with translating ribosomes, transferring the nascent chain between complexes as needed for the specific protein being synthesized. Hence, the presence of ribosomes in our NC-interactomes is not unusual. However, the findings here demonstrate a clear difference in ribosomal protein enrichment in selenos’s NC-interactomes depending on the presence and accessibility of its IDS (Fig. S14). We considered the possibility that this enrichment could be due to contaminants or to stalled synthesis. However, the ribosomal proteins appear to be unique interactors because they were present in the V5-selenos NC-interactome but were greatly diminished or even absent in the V5-selenos 1-133 NC-interactome, even though the two data sets used the same tag position and identical sample-preparation workflow. Furthermore, the impact of the presence and accessibility of selenos’s C-terminus on the interactomes is not limited to ribosomal proteins, as can be seen also for ribosomal biogenesis factors (Fig. S15) and translation initiation factors (Fig. S13).

Ribosomes are enriched at low levels and may therefore appear in the selenos interactome due to associations with membrane protein biogenesis complexes, and not direct interactions with selenos. However, ribosome-nascent chain-associated proteins such as EMC2, CCDC47, Sec61, and Sec63 did not show a pronounced preference for binding V5-selenos over selenos-V5 in NC-interactomes. Hence, it appears unlikely that a selenos protein partner that does not display a preferential binding to V5-selenos is solely responsible for this effect.

To confirm interactions with ribosomes, we used RPS19 as a ribosomal marker in immunoprecipitation assays. We have validated the dependence of selenos interactions with ribosomes on the availability of the redox loop using immunoprecipitation with HA-tagged selenos at either the N- or C-terminus, showing reduced interaction when HA or V5 tags are placed at the C-terminus (Fig. S16A). Hence the observation is independent of the nature of the affinity tag. Additionally, RPS19 was absent in selenos-V5 eluates, and only weak binding was observed for selenos variants lacking the redox loop. These findings are consistent with the NC-interactome results. Variants predicted to adopt an “open” loop conformation – V5-selenos C174A and V5-selenos U188A – interacted with ribosomes similarly to V5-selenos. Moreover, the interaction was not affected by the P178,183S substitutions (Fig. S16B).

In addition to the comparison between the NC-interactomes of V5-selenos and selenos-V5, and a direct comparison of V5-selenos versus selenos-V5, we also analyzed the two full-length selenos NC-interactomes as a grouped dataset (Table S6). This grouped analysis compares protein abundance across both full-length selenos datasets simultaneously against the control of empty vector. Therefore, proteins with different enrichment levels or reproducibility between V5-selenos and selenos-V5 are expected to yield higher P values and are less likely to be retained as significant interactors. Thus, this analysis validates the identification of interactors whose enrichment was not affected by affinity-tag placement (Fig. S17). As expected, ribosomal proteins were largely absent from the grouped analysis because their enrichment differed between the two full-length selenos constructs. This result is consistent with the requirement for an accessible C-terminal redox loop in translation-related interactions, as confirmed by Western blot analysis (Fig. S16).

### The redox loop interacts with membrane protein biogenesis complexes and translation-associated proteins

The R180DiZPK SC-interactome contains a crosslinker positioned within the redox loop, which enables a more refined characterization of interaction partners that directly engage the loop. As noted above, SC-interactome experiments required the use of a C-terminal affinity tag to selectively purify selenos variants containing DiZPK. The presence of this C-terminal affinity may influence interactions occurring within the C-terminal region. Despite this limitation, the R180DiZPK dataset is consistent with the interaction patterns identified in the complementary analyses described above. Specifically, the R180DiZPK SC-interactome is enriched with proteins involved in membrane protein biogenesis, ribosomal components, and RNA-binding proteins (Fig. 5E). This enrichment supports a direct role for the redox loop in engaging factors associated with the translational machinery and insertion of membrane proteins into the ER membrane. In addition, metabolic and redox-active enzymes were detected among the interactors crosslinked at R180. These proteins may be functionally linked to the enzymatic activity of endogenous selenos due to the presence of Sec188. Moreover, cytoskeletal proteins are also enriched in the R180DiZPK SC-interactome (Fig. 5E). This observation aligns with previous reports that selenos functions as a molecular bridge between the ER and microtubules, an interaction that depends on the intrinsically disordered segment (IDS)^33^. Collectively, these data reinforce the view that the redox loop functions as a multifunctional interaction hub that integrates membrane protein insertion, metabolism, redox regulation, cytoskeletal organization, and translational pathways.

## Discussion

We identified selenos’s protein partners using three complementary proteomic approaches (Fig. 1). Firstly, we used AP-MS-DIA, placing the affinity tag at different termini and employing a shorter variant lacking the IDS and its redox loop. This method provided information about tightly bound proteins, their complexes, as well as the changes between different selenos variants and protein partners that require the IDS for strong binding. Secondly, we applied global lysine crosslinking to stabilize interactions between selenos and nearby proteins. This allowed proteins with weaker binding affinities to be detected and made it possible to capture an enriched set of proteins that extends beyond those available by the NC-interactomes alone. When the structures of complexes were available, the crosslinked interactomes enabled us to map which proteins in the specific complex are most likely to interact with selenos by correlating knowledge of their surface lysines with the observations from the crosslinking experiment. Thirdly, we obtained site-specific information about protein partners, by recording interactomes using photo-activatable, site-specific crosslinkers introduced into selenos to capture interactions in live cells. Because crosslinking was carried out *in vivo*, it confirms that these interactions do indeed occur in the living cell. It also allows us to identify transient interactions between selenos and proteins with weaker binding affinities, such as certain redox enzymes. In addition, we were able to determine whether interactions occur in or near the membrane or near the redox loop.

Using this combination of global and site-specific crosslinking, we found that selenos resides *in vivo* near complexes involved in membrane protein biogenesis—the incorporation of newly synthesized proteins into membranes—and their associated quality control pathways (Figs. 2-4). These protein machineries dynamically interact with each other^60^ as the ribosome-nascent chain is moved between them to coordinate the insertion, folding, assembly, and modification of membrane proteins. Selenos’s key partners in this pathway include derlin-1, EMC, OST, SND, and the multipass Sec61 translocon. Our data support multiple contact points for selenos and these biogenesis complexes. Its transmembrane segment is anchored near the OST DAD1/OST3A/B interface. In the EMC complex, its transmembrane segment is placed near EMC3/EMC7 and most likely VDAC, while its cytoplasmic helix contacts EMC2. Selenos can also be placed near TMCO1 and CCDC47 within the multipass translocon (MPT) complex and near the TMEM109 (and possibly the related BRI3BP) and CCDC47 in the SND complex. Tracking the enrichment of relevant complexes in samples crosslinked *in vivo* confirmed that these interactions occur within cells, underscoring their physiological relevance. These complexes are highly likely to be direct interactors of selenos due to their high enrichment in both NC- and crosslinked interactomes, whereas other associated complexes with low enrichment are likely to be pulled down through associations with the OSTA/B, MPT, EMC, and similarly directly bound complexes. Examples of such indirectly associated complexes are the TRAP, SPC, and SRP shown in figure 2. Overall, these findings indicate that selenos is linked to co-translational insertion and post-translational quality control within the ER membrane. While the role of selenos in the EMC, SND, multipass translocon, OST, and other membrane protein biosynthesis pathways is unknown, it is worthwhile noting that several are known to interact with derlin-1. Since selenos is a derlin-1 partner, perhaps it acts as an adapter protein to link derlin-1 to these complexes, especially under cellular stress, to enhance quality control of newly inserted membrane proteins.

In contrast to the high degree of selenos crosslinking to membrane protein biogenesis complexes, the number and enrichment levels of ERAD components were low in the different interactomes, and the degree of crosslinking was also low. Hence, at least in the absence of cellular stress or high levels of unfolded proteins, selenos is not strongly associated with ERAD complexes (Fig. 3 and S7). Indeed, the literature about selenos contributions to ERAD is sometimes conflicting^9^. However, derlin-1 appears to be a common denominator in many protein complexes that contain selenos, such as the OST, EMC, and trafficking complexes. Derlin-1 levels are higher in the cell than those of selenos based on PaxDb 5.0 (Protein Abundance Database)^147^, and it has been reported to reside in complexes with and without selenos^148^. Thus, selenos could couple derlin-1 to specific complexes and to the ATPase p97, regulate access to derlin-1, or control its dislocation activity. Derlin complexes are known to be dynamic and change composition under cellular stress^149^. Here, we suggest that selenos is the adapter coupling derlin-1 to specific complexes.

Furthermore, mitochondrial membrane proteins are among the enriched proteins in the selenos interactome and are also detected in crosslinking interactomes. Among the most enriched mitochondrial proteins are the VDAC ion channels, which regulate calcium influx and the transport of other metabolites and contribute to stress responses. VDAC is coupled to the EMC at MAMs. Multiple enriched proteins that are also crosslinked are in the immediate vicinity of the VDACs. In addition, complex IV (Fig. 4C) is present in crosslinked interactomes. The finding that selenos is linked to mitochondrial processes and potentially to regulation of OXPHOS is supported by reports in the literature that calcium homeostasis^19^ and mitochondrial function^137^ are disrupted upon selenos depletion. Indeed, the ER and mitochondria’s functions in stress responses are intertwined^150^. However, our studies here narrow down the involvement to three likely routes: i) Selenos engagement with EMC/VDAC complexes (Fig. 2); ii) Selenos presence in the PHB1/PHB2 complex and putatively assembly and degradation of mitochondrial electron transport chain complexes (Fig. 4). iii) Association of selenos with Sarco-Endoplasmic Reticulum Ca²⁺-ATPase (SERCA), a calcium ion pump that transports Ca²⁺ from the cytosol to the ER^151^, and is enriched in NC-interactomes (Fig. S7E).

Selenos is notable because it is one of only 25 human proteins that incorporate the reactive amino acid Sec, yet the physiological role of this Sec residue remains unknown^9^. The redox loop of selenos, in which the Sec resides, has several unique features that render it suitable for cellular control. Firstly, the low redox potential of the selenenylsulfide bond between the loop’s Sec and Cys^23^ means that it can exist in the cytoplasm despite its reducing environment. It can be broken down only by a select few enzymes. For example, we have previously shown that thioredoxin reductase 1 can reduce selenos *in vitro*^23^. Secondly, the conformation of the redox loop itself is confined by the presence of two prolines and positively charged residues. The length of the spacing residues and the loop conformation control the encounter rate between the Sec and its partnering Cys, as well as the reformation of the selenylsulfide bond. So, even though the number of residues separating the Cys and Sec in the redox loop is unusually large for a selenoprotein, we have shown that the rate of selenylsulfide reformation *in vitro* is extremely fast^22^. Here, we were able to provide evidence for processes that require the redox loop.

The complex that most clearly relies on the redox loop to mediate interactions is the elongator subcomplex ELP1/ELP2/ELP3 (Fig. 5), which catalyzes tRNA modifications and is important for efficient and accurate translation^152^. Consistent with this observation, selenos levels are known to increase under cellular stress^9^, a condition in which alterations in tRNA modifications influence translation^153,154^. Our proteomics data indicate that recruitment of the elongator complex requires an exposed and accessible redox loop, a finding that we corroborated by immunoprecipitation experiments. Crosslinking proteomics and immunoprecipitation further revealed that this observation also applies to Sec63, a membrane protein biogenesis factor that appears at moderate levels in NC-interactomes but was specifically captured in the R180DiZPK interactome. Some of the translation-related partners, such as eIF5A, are only present in crosslinked interactomes. Ribosome biogenesis factors and ribosomes were significantly more abundant in the V5-selenos NC-interactome than in those of V5-selenos 1-133 or selenos-V5 NC-interactomes (Fig. 5), suggesting the importance of the presence and accessibility of the redox loop for the interaction. Regrettably, it was only technically possible to utilize selenos variants with a C-terminal tag for our site-specific interactomes, which is a technical limitation of the method for studying the translation-related proteins. A role for selenos in the regulation of translation is consistent with the broader observation that selenoprotein expression is elevated under stress, reflecting its role in stress management^155^.

In conclusion, our findings extend what is known about the role of selenos to encompass a broader role in membrane protein biogenesis and in translation. They indicate engagement with multiple machineries that link translation, membrane protein insertion, quality control, metabolism, and trafficking at the ER membrane. Additionally, there is a link between selenos and mitochondrial function, which appears to be coupled to membrane protein biogenesis complexes at the MAM but also encompasses the regulation of mitochondrial respiratory complexes that carry out oxidative phosphorylation. We provide evidence that processes related to translation preferentially require selenos’s redox loop, as validated by crosslinking experiments capturing interactions in live cells and immunoprecipitation. An accessible and conformationally competent redox loop is required for association with ribosomes, ribosome biogenesis factors, and key regulators of translation, including the elongator complex. These findings establish a direct mechanistic link between Sec-containing redox chemistry in selenos and translational regulation, supporting a model in which the redox loop functions as a regulated interaction module that couples selenos to cotranslational protein biogenesis.

## Materials and Methods

### Materials

HEK293T cells were from ATCC (CRL-3216). Dulbecco’s Modified Eagle’s Medium (DMEM) from Corning (#10-013-CV) was used for culturing human cells. The Q5™ Site-Directed Mutagenesis Kit (#E0552S) was from New England Biolabs (NEB). Transfection was done using Polyjet reagent (SignaGen Laboratories #SL100688) and polyethyleneimine (PEI, 25 kDa, Kyfora Bio #23966). Igepal CA-630 was obtained from Sigma-Aldrich #I8896. Primers were synthesized by Sigma-Aldrich. The NucleoSpin mini kit for plasmid DNA was obtained from Machery-Nagel # 740588. The ZymoPURE II Plasmid Midiprep Kit was purchased from Zymo Research #D4200. Dulbecco’s Modified Eagle’s Medium (DMEM) was obtained from Corning #10-013-CV, fetal bovine serum (FBS) from GenClone #25-550, and penicillin-streptomycin from HyClone #SV30010. Protease inhibitor cocktail (Genesee Scientific #18-430) and phosphatase inhibitor cocktail III (Genesee Scientific #18-451B) were used in cell lysates. V5-trap magnetic agarose (ChromoTek #v5tma) was used for affinity purification. For the list of antibodies and plasmids used in this study, see Table S7.

### Protein expression plasmids

The *Homo sapiens* SELENOS gene (GenBank accession no. GI: 45439348) was cloned into a modified pcDNA5/FRT/TO vector (Addgene #105769) following procedures described in Szczesny et al.^156^. The resulting construct included the native SELENOS 3′ untranslated region (3′-UTR), including the endogenous SECIS element. To achieve homogeneous expression, Sec188 was mutated to Cys. A V5 affinity tag (GKPIPNPLLGLDST) was inserted at either the N-or C-terminus of selenos.

### Generation of stably expressing cell lines

Flp-In T-REx 293 cells (Thermo Fisher Scientific) were used for the generation of stable cell lines using the method explained in reference^67^. Briefly, 500,000 cells were seeded into 6-well plates in DMEM containing 10% (v/v) FBS and 1% (v/v) penicillin-streptomycin. Cells were incubated in a humidified incubator at 37 °C, 5% CO_2_. The next day, cells were transfected with 1 μg of the pOG44 Flp-recombinase expression vector (Thermo Fisher Scientific #V600520), 0.3 μg of the expression vector, and 4 μL of Polyjet according to the manufacturer’s instructions. The next day, the medium was changed to selection media (DMEM, 10% (v/v) FBS, 1% (v/v) penicillin-streptomycin, 50 μg/mL hygromycin B (GoldBio #H-270-1), and 10 μg/mL blasticidin S (GoldBio #B-800). The selection medium was changed every 3 days until colonies formed. As a selection control, only pOG44 was transfected into the cells and grown in the selection medium. Cells with an insert grew in the selection medium, while the cells transfected only with pOG44 did not survive. Addgene plasmid #105769 was used to generate stable cell lines for AP-MS-DIA controls under identical antibiotic selection conditions, without protein overexpression.

### Cell culture for mass spectrometry

Stably expressing cell lines containing V5-tagged selenos and cell lines containing empty vector (five repeats each) were grown in DMEM containing 10% (v/v) FBS and 1% (v/v) penicillin-streptomycin. Cells were grown in two 15 cm dishes and incubated in a humidified incubator at 37 °C, 5% CO_2_. Expression of the gene in Flp-In T-Rex 293 derived stable cell lines were induced by adding 100 ng/mL doxycycline to the medium after they reached about 70% confluency. After 48 h, the cells were washed twice with cold PBS (137 mM NaCl, 2.7 mM KCl, 10 mM Na_2_HPO_4_, and 1.8 mM KH_2_PO_4_), scraped and the pellets were kept at -80 °C until further processing.

### Site-specific crosslinker incorporation and crosslinking

HEK293T cells were grown in DMEM containing 10% FBS (v/v) and 1% (v/v) penicillin-streptomycin. Cells were grown in two 15 cm dishes and were incubated in a humidified incubator at 37 °C, 5% CO_2_. A 200 mM (S)-2-amino-6-(3-(3-(3-methyl-3H-diazirin-3-yl)propyl)ureido)hexanoic acid (DiZPK) (aablocks #AA009CEV) stock was made in 200 mM HCl and added to the medium to a final concentration of 0.2 mM, and the pH was neutralized with an equal volume of 200 mM NaOH. After cells reached 70% confluency, the medium was changed to the DiZPK-containing medium. Cells were transfected with equal amounts of plasmid encoding for the desired V5 variant and the vector containing the aminoacyl-tRNA synthetase/tRNA pair (Addgene plasmid #91706) with PolyJet (SignaGen Laboratories # SL100688) following the manufacturer’s instructions. After 24 h, the medium was replaced with growth medium containing the UAA. After another 24 h, the cells were washed twice with cold PBS (137 mM NaCl, 2.7 mM KCl, 10 mM Na_2_HPO_4_, and 1.8 mM KH_2_PO_4_). The 15 cm plates with cells submerged in 5 mL PBS were kept on ice. Crosslinking was performed by irradiating the cells at 365 nm for 20 min using a Stratalinker 2400 UV Crosslinker (Stratagene), with plates positioned 9 cm below the light source. The power at 365 nm was measured to be 2.0 mW. The control dishes were kept at 4 °C for the same amount of time. Cells were then scraped, and the pellets were kept at -80 °C until pull-down and further processing.

### Global lysine crosslinking

Stably expressing cells containing V5-selenos U188C (three repeats) were grown in DMEM containing 10% (v/v) FBS and 1% (v/v) penicillin-streptomycin. Cells were grown in two 15 cm dishes and were incubated at 37 °C, 5% CO_2_. Expression of the gene in Flp-In system cell lines was driven by inducible promoters. The cells were induced by adding 100 ng/mL doxycycline to the medium after they reached about 70% confluency. After another 48 h, the cells were washed twice with cold PBS (137 mM NaCl, 2.7 mM KCl, 10 mM Na_2_HPO_4_, and 1.8 mM KH_2_PO_4_). A 200 mM Di(N-succinimidyl) 3,3’-dithiodipropionate (DSP) (AK Scientific # 6600AH) stock solution in dimethyl sulfoxide (DMSO) was prepared fresh and then diluted to 0.8 mM in PBS. The DSP solution was added to the cells and incubated at 37 °C for 20 min. For the control condition, an equal amount of DMSO was added to PBS. After removing the crosslinking solution, the crosslinking reaction was quenched with 5 mL of 25 mM Tris-HCl (pH 7.4) at room temperature for 10 min, then the cells were scraped and the pellets were kept at -80 °C until pull-down and further processing.

### Affinity purification for mass spectrometry

Cells were thawed on ice for 20 min and lysed with 2 mL of cold lysis buffer (10 mM Tris-HCl, pH 7.4, 150 mM NaCl, 0.5% (v/v) Igepal CA-630, 0.5 mM EDTA) supplemented with 10 μL/mL protease inhibitor and phosphatase inhibitor cocktail. Cells were lysed with 6 freeze-thaw cycles and centrifuged at 13,000 x g for 20 min at 4 °C to pellet cell debris. The supernatant was collected and mixed with 33 μL of V5-trap magnetic agarose. After 4 h of incubation with mixing at 4 °C, beads were washed once with 1 mL of cold wash buffer (10 mM Tris-HCl, pH 7.4, 150 mM NaCl, 0.05% (v/v) Igepal CA-630, 0.5 mM EDTA) and twice with 1 mL of cold 100 mM ammonium hydrogen carbonate buffer. Approximately 10% of the beads were set aside for immunoblot analysis, and the remaining beads were stored at −80 °C until digestion. As noted above, the controls consisted of stable cell lines generated with an empty vector and were subjected to the same growth conditions and sample preparation procedures.

### Western blot

Transfer from SDS-PAGE gels was carried out using a power blotter semi-dry system (Thermo Fisher Scientific). The membrane was blocked with 5% (w/v) BSA in TBST (Tris-buffered saline, 0.1% (v/v) Tween 20) for 1 h. The membrane was then incubated with the primary antibody for 1 h, washed three times, incubated with horseradish peroxidase (HRP)-conjugated secondary antibody for 1 h, and washed three times. The western blot was visualized using a chemiluminescent HRP substrate kit (Genesee # 20-300B). Images were taken using a FluorChem R System (biotechne).

### AP-MS-DIA and crosslinking protein digestion and LC-MS/MS

Protein samples were processed using the Filter-Aided Sample Preparation (FASP) method ^157^ over two days. On Day 1, on-bead protein samples were first resuspended in 200 µL of 100 mM triethylammonium bicarbonate (TEAB) and reduced by adding 10 µL of 200 mM tris(2-carboxyethyl)phosphine hydrochloride (TCEP), followed by incubation at 37 °C for 30 min. Meanwhile, FASP filter units were rinsed with 400 µL of HPLC-grade water and centrifuged at 14,000 × g for 10 min. Reduced samples were loaded onto the filters and centrifuged for 20 min at the same speed. For alkylation, 100 µL of 30 mM iodoacetamide (IAA) in TEAB was added, gently vortexed, and incubated in the dark for 20 min without mixing. The samples were then centrifuged again, followed by three washes with 300 µL of 100 mM TEAB, each with centrifugation at 14,000 x g for 20 min. Filter units were transferred to fresh collection tubes, and 100 µL of 100 mM TEAB buffer was added.

Proteins were digested by adding 10 µL of trypsin solution to each sample, achieving an enzyme-to-protein ratio of 1:50. The trypsin solution was prepared by reconstituting 20 µg of sequencing-grade modified porcine trypsin (Promega) in 100 µL of 0.01% (v/v) formic acid containing 50 mM calcium ions. Samples were gently vortexed and incubated in a humid chamber at 47 °C for 4-18 h. On day 2, peptides were recovered by centrifugation at 14,000 × g for 10 min, followed by an additional wash with 80 µL of 100 mM TEAB and a second centrifugation. Samples were then acidified to a final concentration of 0.5% formic acid and 0.1% trifluoroacetic acid (TFA) by adding 20 µL of a 5% (v/v) formic acid, 1% (v/v) TFA solution. Peptide mixtures were desalted using Sep-Pak plates according to the manufacturer’s standard protocol.

Tryptic peptides were then separated by reverse phase XSelect CSH C18 2.5 um resin (Waters) on an in-line 150 x 0.075 mm column using an UltiMate 3000 RSLCnano system (Thermo Scientific). Peptides were eluted using a 60 min gradient from 98:2 to 65:35 buffer A:B ratio. Eluted peptides were ionized by electrospray (2.4 kV) through a heated capillary (275 °C) followed by data collection on an Orbitrap Exploris 480 mass spectrometer (Thermo Scientific).

Precursor spectra were acquired with a scan from 385-1015 Th at a resolution set to 60,000 with 100% AGC, max time of 50 ms, and an RF parameter at 40%. DIA was configured on the Orbitrap 480 to acquire 40 x 15 Th isolation windows, normalized AGC target 500%, maximum injection time 40 ms. A second DIA was acquired in a staggered window (15 Th) pattern with optimized window placements from 390-990 Th. Buffer A = 0.1% (v/v) formic acid, 0.5% (v/v) acetonitrile, Buffer B = 0.1% (v/v) formic acid, 99.9% (v/v) acetonitrile.

### AP-MS-DIA and crosslinking statistical analysis

Following data acquisition, the data were searched using Spectronaut (Biognosys version 20.1) against the UniProt *Homo sapiens* database (6^th^ version of 2024) using the directDIA method with an identification precursor and protein q-value cutoff of 1%, generate decoys set to true, the protein inference workflow set to maxLFQ, inference algorithm set to IDPicker, quantity level set to MS2, cross-run normalization set to false, and the protein grouping quantification set to mean peptide and precursor quantity. Each set of bait samples was filtered for proteins identified with at least two peptides. Protein MS2 intensity values were assessed for quality using ProteiNorm^158^. For AP-MS-DIA, only proteins present in at least four out of five replicates, and for crosslinking proteins present in two out of three replicates, were kept for analysis. The data were normalized using log_2_ intensities with the quantile method, and missing values were imputed with the BPCA method^159^ using the NAguideR R package^160^. The proteins were analyzed using proteoDA^161^ to perform statistical analysis using Linear Models for Microarray Data (limma) with empirical Bayes (eBayes) smoothing to the standard errors^162,163^. For differential enrichment analysis, protein abundances quantified in the NC-interactome were compared with those in empty vector control samples. In the LC-interactome, protein abundances in crosslinked samples were compared with those in the corresponding non-crosslinked (no DSP treatment) samples. In the SC-interactome, protein abundances in UV-irradiated samples were compared with those in samples kept in the dark (no photoactivation). Proteins with an FDR-adjusted p-value < 0.05 and a fold change > 2 were considered significant (Fig. S19). Gene ontology analysis was done for biological processes. Terms were grouped by Metascape^141^, and the representative terms with the highest adjusted p-values were shown.

### Immunoprecipitation for immunoblotting and bait verification

HEK293 cells were grown in DMEM containing 10% (v/v) FBS and 1% penicillin-streptomycin. Cells were incubated at 37 °C, 5% CO_2_. Cells were transfected with the expression plasmid using a 1:3 ratio of polyethylenimine (PEI, 25 kDa, Kyfora Bio #23966). The medium was exchanged after 24 h. After another 24 h, the cells were harvested and washed twice with cold PBS (137 mM NaCl, 2.7 mM KCl, 10 mM Na_2_HPO_4_, and 1.8 mM KH_2_PO_4_). Cells were snap-frozen and kept at - 80 °C until pull-down. Experiments were carried out in three biological replicates.

Cells were thawed on ice for 15 min and lysed with 200 μL of cold lysis buffer (10 mM Tris-HCl, pH 7.4, 150 mM NaCl, 0.5% Igepal CA-630, 0.5 mM EDTA) supplemented with 10 μL/mL protease inhibitor and phosphatase inhibitor cocktails. Cells were lysed with six freeze-thaw cycles and centrifuged at 13,000 x g for 20 min at 4 °C to pellet cell debris. The supernatant was collected and mixed with 3 μL of V5-trap magnetic agarose. After 4 h of incubation with mixing at 4 °C, beads were washed twice with 200 μL of cold wash buffer (10 mM Tris-HCl pH 7.4, 150 mM NaCl, 0.05% (v/v) Igepal CA-630, 0.5 mM EDTA) and once with 200 μL of 10 mM Tris-HCl pH 7.4, 150 mM NaCl, 0.5 mM EDTA. Beads were eluted with 20 μL of 2x Laemmli buffer and boiled at 95 °C for 5 min.

Measurements of band intensities were performed using Fiji, and the intensities were normalized for each membrane. At least three biological replicates were used for each condition.

### Immunocytochemistry and colocalization analysis

Staining followed a modified version of the protocol described in reference^164^. In this procedure, 100,000 Flp-In T-REx 293 cells stably expressing V5-selenos U188C, selenos-V5 U188C, V5-selenos 1-133, or selenos U188C were seeded onto gelatin-coated coverslips (no. 1.5 thickness) in 6-well plates. Induction was achieved using 100 ng/mL doxycycline, with cells incubated for 48 h. Cells were rinsed three times with PBS before being fixed with 4% (w/v) formaldehyde in growth medium at room temperature for 15 min. Following fixation, cells were permeabilized twice with 0.01% (v/v) Triton X-100 in PBS for 3 min each and blocked in 5% BSA in PBS for 1 h. Primary antibodies were then applied at a 1:200 dilution in blocking buffer and incubated overnight at 4 °C. After four PBS washes of 5 min each, cells were incubated with secondary antibodies (1:200 dilution) in the dark at room temperature. Coverslips were mounted onto slides with ProLong Gold Antifade Mountant (Invitrogen #P36934) and sealed after final washes. Samples were allowed to cure overnight, and imaging was performed with a 63x/1.4NA/oil-immersion objective on a Zeiss LSM 880 with Airyscan. Image processing was conducted using ZEN Blue software (Zeiss). At least 20 images from three biological replicates were acquired.

Colocalization between the indicated fluorescence channels was quantified in Fiji using the JACoP plugin by calculating the Manders’ M1 coefficient. Prior to analysis, images were background corrected, and an automatic Otsu threshold was applied independently to each channel to define signal-positive pixels. Manders’ M1 was then computed as the fraction of signal intensity in the V5 (selenos) channel overlapping with the thresholded signal in the calnexin (ER) channel. At least 20 images from three biological replicates were analyzed for an unpaired two-sample Welch’s t-test analysis (over 25 cells in each sample).

## Acknowledgment

We thank the Institutional Development Award (IDeA) National Resource for quantitative proteomics, especially Drs. Province, Mackintosh and Edmondson, for their expert advice and assistance with data acquisition and analysis. This project was funded by the National Science Foundation under grant 2150863 and the National Institute of General Medical Sciences (NIGMS) of the National Institutes of Health under the award number 1R35GM153494 to Sharon Rozovsky at the University of Delaware. It was also supported by NIGMS awards number R01AI186337 and R01AG078925 to John La Cava at The Rockefeller University. Atinuke Odunsi was funded in part by NIGMS Chemistry-Biology Interface fellowship T32-GM133395. Farid Ghelichkhani was partly supported by the University of Delaware Graduate College Doctoral Fellowship Award. Mariia A. Kapitonova was supported in part by an American Heart Association 23IPA1053632. Instrumentation was supported by National Institute of General Medical Sciences Award Number P30GM159572. The generation of HEK293 SELENOS knock-out cell lines was supported by the Institutional Development Award from the National Institute of Health’s National Institute of General Medical Sciences under grant number P20GM103446. Funding for the IDeA National Resource for quantitative proteomics is provided through a grant from the National Institute of General Medical Sciences (R24GM137786). The content is solely the responsibility of the authors and does not necessarily represent the official views of the National Institutes of Health. Fig. 1 was created in part using BioRender.com.

## Author contributions

F. G. acquired and analyzed data

A.O., M.A.K., and E.R. assisted in data collection.

O.G.R., M.H.K., and J.L. assisted with proteomics data analysis

F.G. and S.R. wrote the manuscript, and all authors reviewed and edited it. All authors contributed to fact checking and revisions of the article.

## Data Availability Statement

The MS raw files associated with this study have been deposited in PRIDE/ProteomeXchange (https://proteomecentral.proteomexchange.org) with the accession number PXD075535.

## Supporting Information

### Supplemental Tables

Table S1: AP-MS-DIA output

Table S2: DSP global crosslinking output

Table S3: DiZPK photocrosslinking output

Table S4: Comparison of the AP-MS-DIA interactomes of V5-selenos and V5-selenos 1-133

Table S5: Comparison of the AP-MS-DIA interactomes of V5-selenos and selenos-V5

Table S6: Analysis of V5-selenos and selenos-V5 grouped together compared with empty-vector controls.

Table S7: Antibodies, plasmids and protein sequences

### Supplemental Figures

Figure S1. Short linear motifs (SLiMs) in selenos predicted using the Eukaryotic Linear Motif (ELM) resource.

Figure S2. Selenos expression, Golgi localization, and immunofluorescence analysis of cytoskeletal and er morphological changes.

Figure S3. Western blot analysis of samples used for proteomics.

Figure S4. Reciprocal pull-down of selenos using EMC2 as the bait.

Figure S5. Selenos is not required for preemptive quality control or glycosylation of selected proteins.

Figure S6. Summary of the ER protein biogenesis machinery of the components found in selenos’s interactome in this study.

Figure S7. ERAD components and ATPases in selenos interactomes.

Figure S8. The overlap between selenos and derlins interactomes.

Figure S9. Mitochondrial proteins in the selenos interactome.

Figure S10. ER-mitochondria contact sites are not changed in SELENOS knockout (KO) and HEK293 cells.

Figure S11. Gene Ontology (GO) enrichment analysis of biological processes by Metascape for V5-selenos, selenos-V5 and V5-selenos 1-133.

Figure S12. The interaction between selenos and ELP1 is reduced by a C-terminal affinity tag, regardless of tag identity.

Figure S13. Translation initiation factors identified in the selenos interactomes.

Figure S14. Ribosomal proteins identified in the selenos interactome.

Figure S15. Ribosomal biogenesis factors identified by AP-MS-DIA. Figure S16. Selenos associates with ribosomes.

Figure S17. Comparison of grouped and individual NC-interactome analyses of full-length selenos.

Figure S18. Biological replicates of the experiments shown in Figs. 5D and S16B.

Figure S19. Workflow of the statistical analysis applied to the MS datasets.

