## Supplemental information for "Selenoprotein S plays a role in translation and membrane protein biogenesis"

### Supplemental Tables:

Table S1: AP-MS-DIA output
Table S2: DSP global crosslinking output
Table S3: DiZPK photocrosslinking output
Table S4: Comparison of the AP-MS-DIA interactomes of V5-selenos and V5-selenos 1–133
Table S5: Comparison of the AP-MS-DIA interactomes of V5-selenos and selenos-V5
Table S6 Analysis of V5-selenos and selenos-V5 grouped together compared with empty-vector controls.
Table S7: Antibodies, plasmids and protein sequences

### Supplemental Figures:

Figure S1. Short linear motifs (SLiMs) in selenos predicted using the Eukaryotic Linear Motif (ELM) resource.
Figure S2. Selenos expression, Golgi localization, and immunofluorescence analysis of cytoskeletal and er morphological changes.
Figure S3. Western blot analysis of samples used for proteomics.
Figure S4. Reciprocal pull-down of selenos using EMC2 as the bait.
Figure S5. Selenos is not required for preemptive quality control or glycosylation of selected proteins.
Figure S6. Summary of the ER protein biogenesis machinery of the components found in selenos's interactome in this study.
Figure S7. ERAD components and ATPases in selenos interactomes.
Figure S8. The overlap between selenos and derlins interactomes.
Figure S9. Mitochondrial proteins in the selenos interactome.
Figure S10. ER-mitochondria contact sites are not changed in SELENOS knockout (KO) and HEK293 cells.
Figure S11. Gene Ontology (GO) enrichment analysis of biological processes by Metascape for V5-selenos, selenos-V5 and V5-selenos 1-133.
Figure S12. The interaction between selenos and ELP1 is reduced by a C-terminal affinity tag, regardless of tag identity.
Figure S13. Translation initiation factors identified in the selenos interactomes.

Figure S14. Ribosomal proteins identified in the selenos interactome.
Figure S15. Ribosomal biogenesis factors identified by AP-MS-DIA.
Figure S16. Selenos associates with ribosomes.
Figure S17. Comparison of grouped and individual NC-interactome analyses of full-length selenos.
Figure S18. Biological replicates of the experiments shown in Figs. 5D and S16B.
Figure S19. Workflow of the statistical analysis applied to the MS datasets.

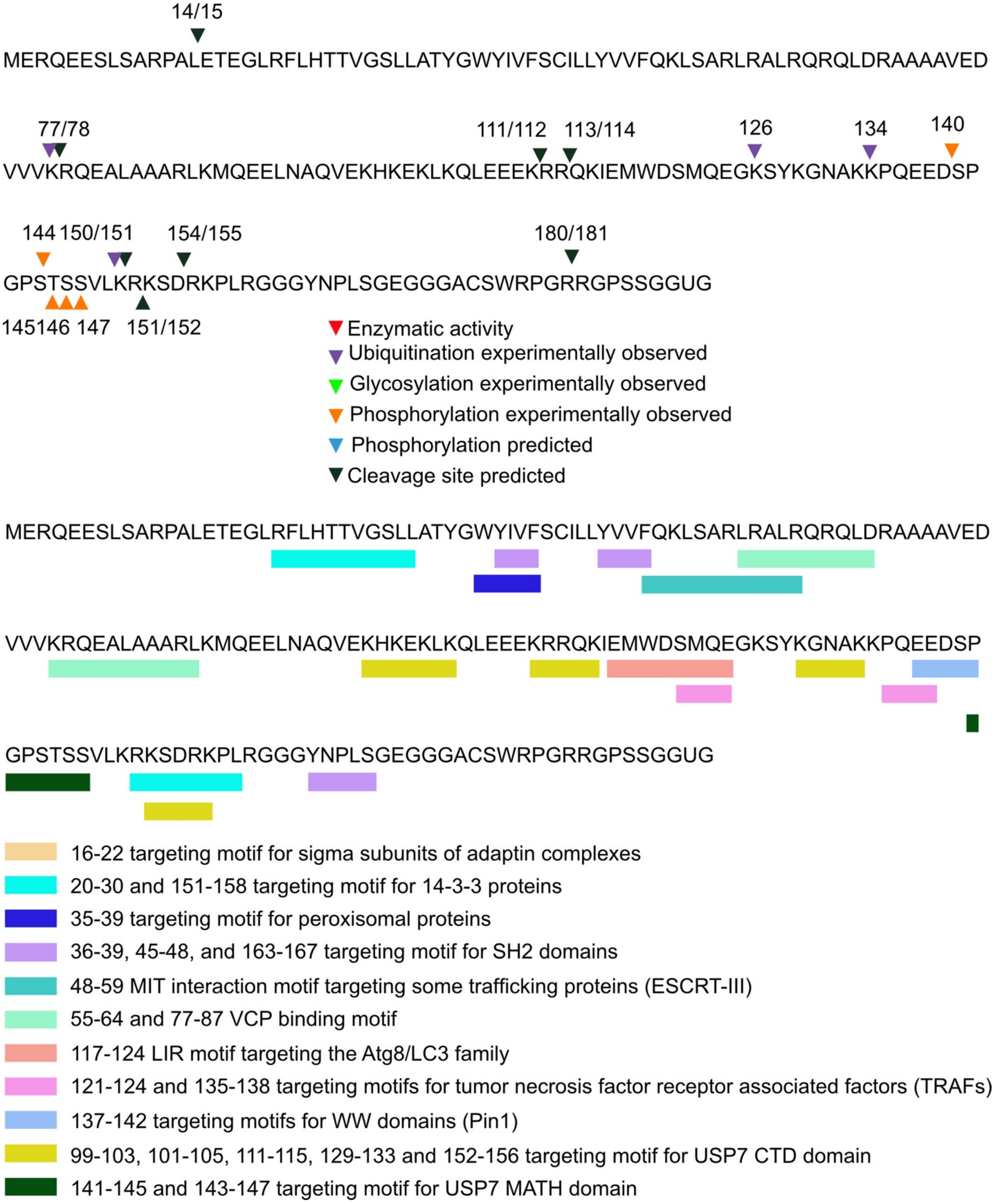


Figure S1. Short linear motifs (SLiMs) in selenos predicted using the Eukaryotic Linear Motif (ELM) resource[1].

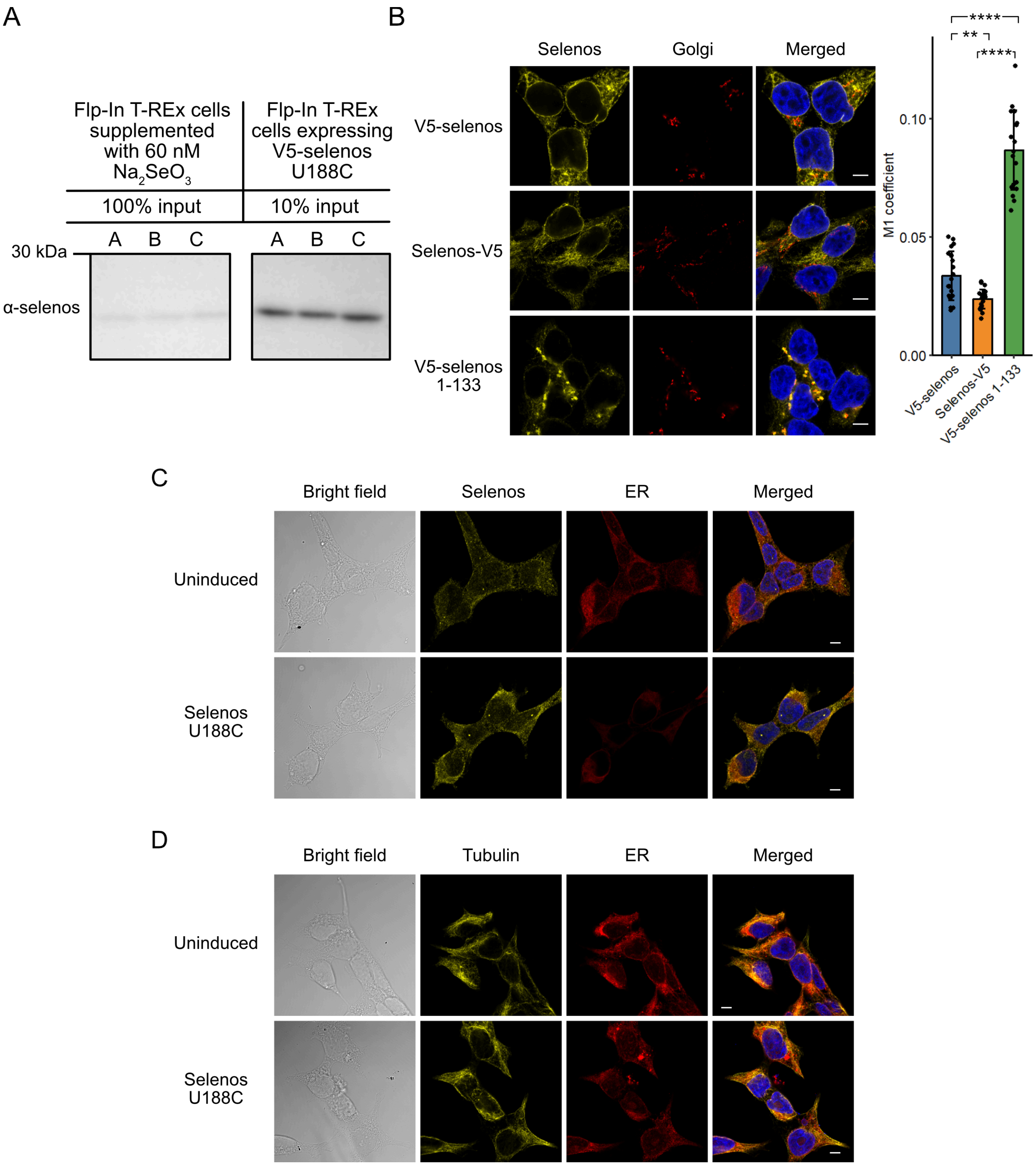


Figure S2. Selenos expression, Golgi localization, and immunofluorescence analysis of cytoskeletal and ER morphological changes. A) Comparison of overexpressed selenos protein levels with endogenous selenos levels. Under the experimental conditions used for AP-MS-DIA, induction of V5-selenos U188C with 100 ng/mL doxycycline resulted in protein levels approximately 58-fold higher than the endogenous selenos levels with selenium supplementation. Considering the presence of endogenous selenos in Flp-In T-REx 293 cells, this corresponds to a total selenos level comparable to endogenous levels. Sample concentrations were normalized to a total protein content of the lysates, and appropriate dilutions were applied to prevent signal overexposure. Measurements were performed in three biological replicates, and intensity quantification was carried out using FIJI. It is noteworthy that selenos levels have been reported to increase approximately fourfold in response to ER stress[2]. All conditions were blotted on the same membrane to enable direct comparison of protein yield. B) Immunofluorescence microscopy of selenos variants (V5 antibody) (yellow), cis-Golgi marker GM130 (red), and nuclei (blue). Scale bar, 5 µm. The M1 coefficient was used to quantify selenos localization to the Golgi. Error bars show mean ± SD. At least 20 images from 3 biological replicates were analyzed. T-test ****$p < 0.0001$; **$0.01 < p < 0.001$; ns, not significant. C) Flp-In T-REx 293 cells expressing selenos U188C were used to assess the impact of overexpression on selenos localization. Cells were either uninduced or induced with 100 ng/mL doxycycline, matching the AP-MS experimental conditions. D) Tubulin and calnexin were immunostained in both uninduced and induced cells. The images show normal ER and cytoskeletal morphology under both conditions, indicating that the expression levels used do not induce cellular stress and do not affect selenos localization.

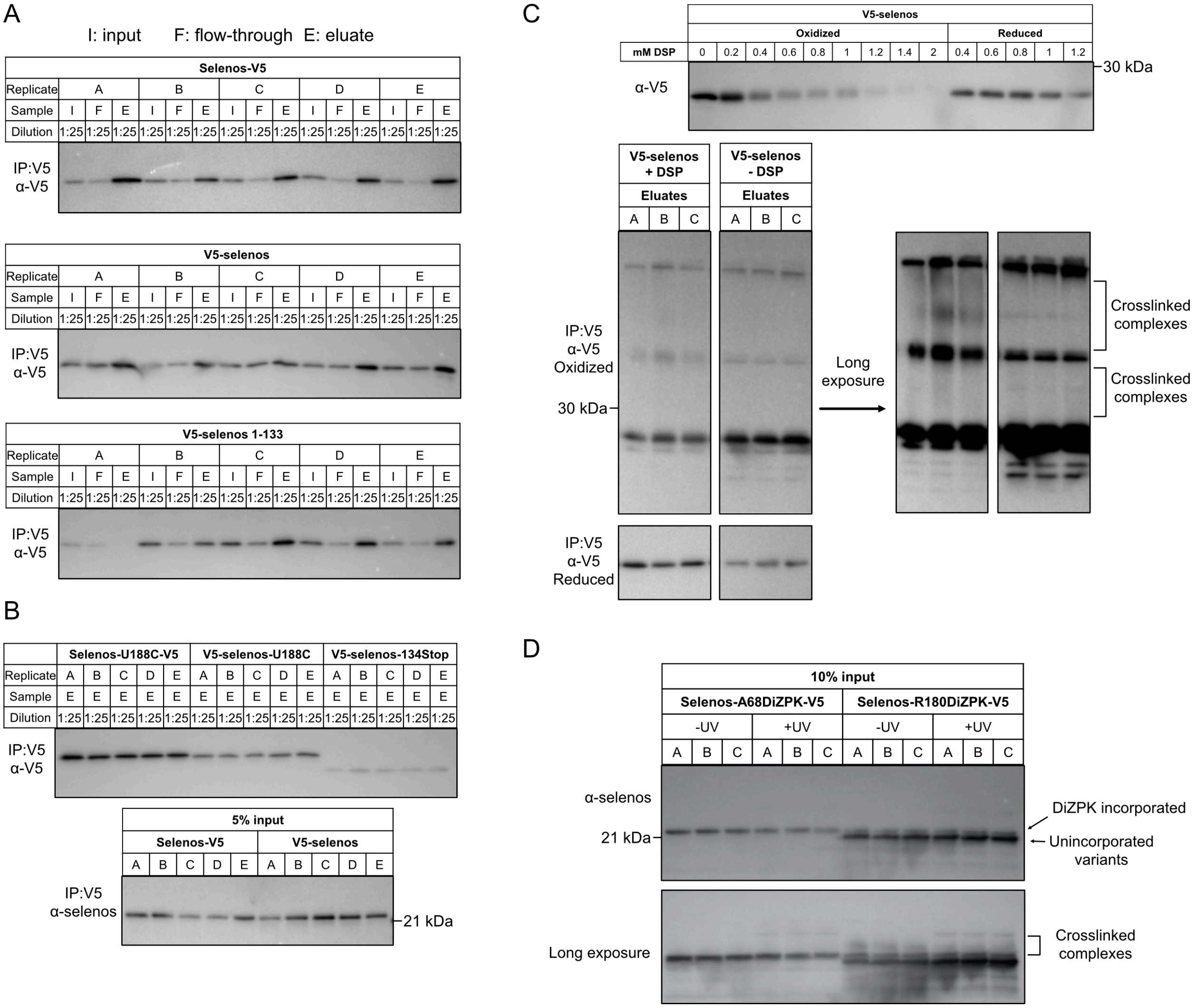


Figure S3. Western blot analysis of samples used for proteomics. A) Western blots of AP-MS-DIA samples. Samples were diluted to ensure detection within the linear range of the chemiluminescence signal. B) All conditions were blotted on the same membrane to enable direct comparison of protein yield. For this specific V5 antibody, binding to N-terminal V5 tags is weaker, resulting in a lower signal. The membrane probed with anti-selenos antibody shows that bait protein levels are approximately similar between V5-selenos and selenos-V5. The truncated variant, V5-selenos 1–133, is expressed at ~40% of the full-length V5-selenos level, despite using the same expression system, suggesting reduced stability. C) Optimization of DSP crosslinker concentration was performed using Flp-In T-REx 293 cells expressing V5-selenos (top panel). Increasing DSP concentration led to the disappearance of the V5-selenos band, indicating the formation of higher-order complexes. Upon reduction, these complexes were disrupted. A concentration of 0.8 mM DSP was selected to balance under- and over-crosslinking. The bottom panel shows eluates after IP. Under oxidizing conditions, selenos forms multimers that are detectable even in samples without DSP. Longer exposure reveals high-molecular-weight complexes formed after crosslinking. D) Whole-cell lysates from cells expressing selenos-V5 containing A68DiZPK or R180DiZPK were analyzed prior to immunoprecipitation to assess yield and crosslinking efficiency. The unincorporated variant of A68DiZPK is not detectable, likely due to instability in cells or its small size limiting detection by western blot.

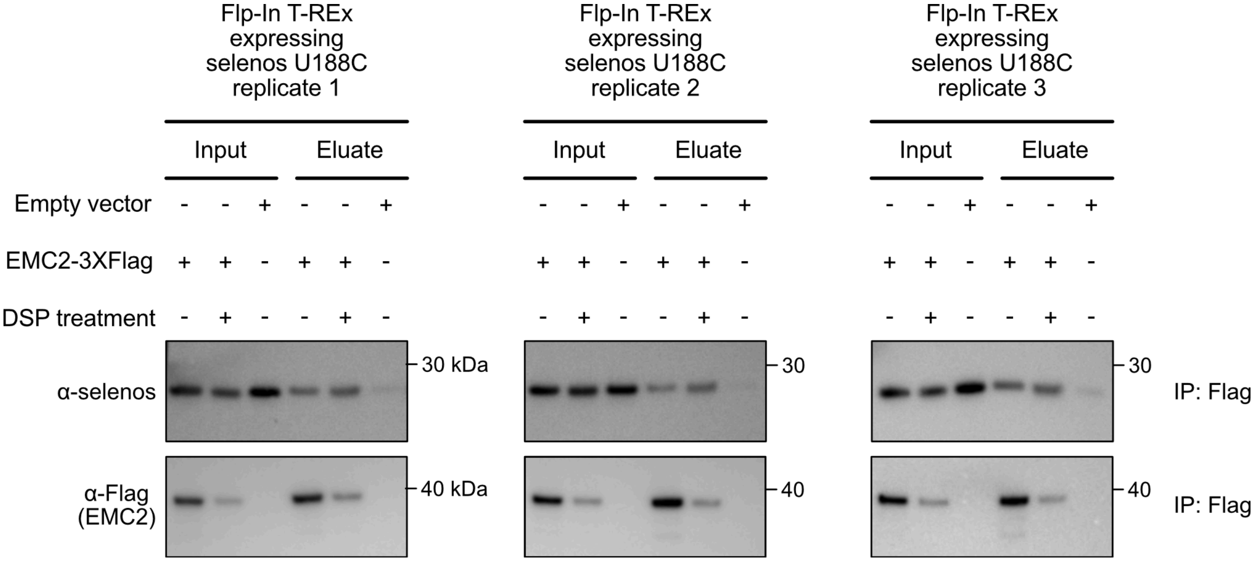


Figure S4. Reciprocal pull-down of selenos using EMC2 as the bait. EMC2 carrying a C-terminal 3XFlag tag was transiently expressed in Flp-In T-REx 293 cells expressing selenos U188C (tagless selenos). Selenos and EMC2 expression was induced with 100 ng/mL doxycycline for 48 h. Immunoprecipitation was performed using anti-Flag magnetic beads (Pierce, #A36797).

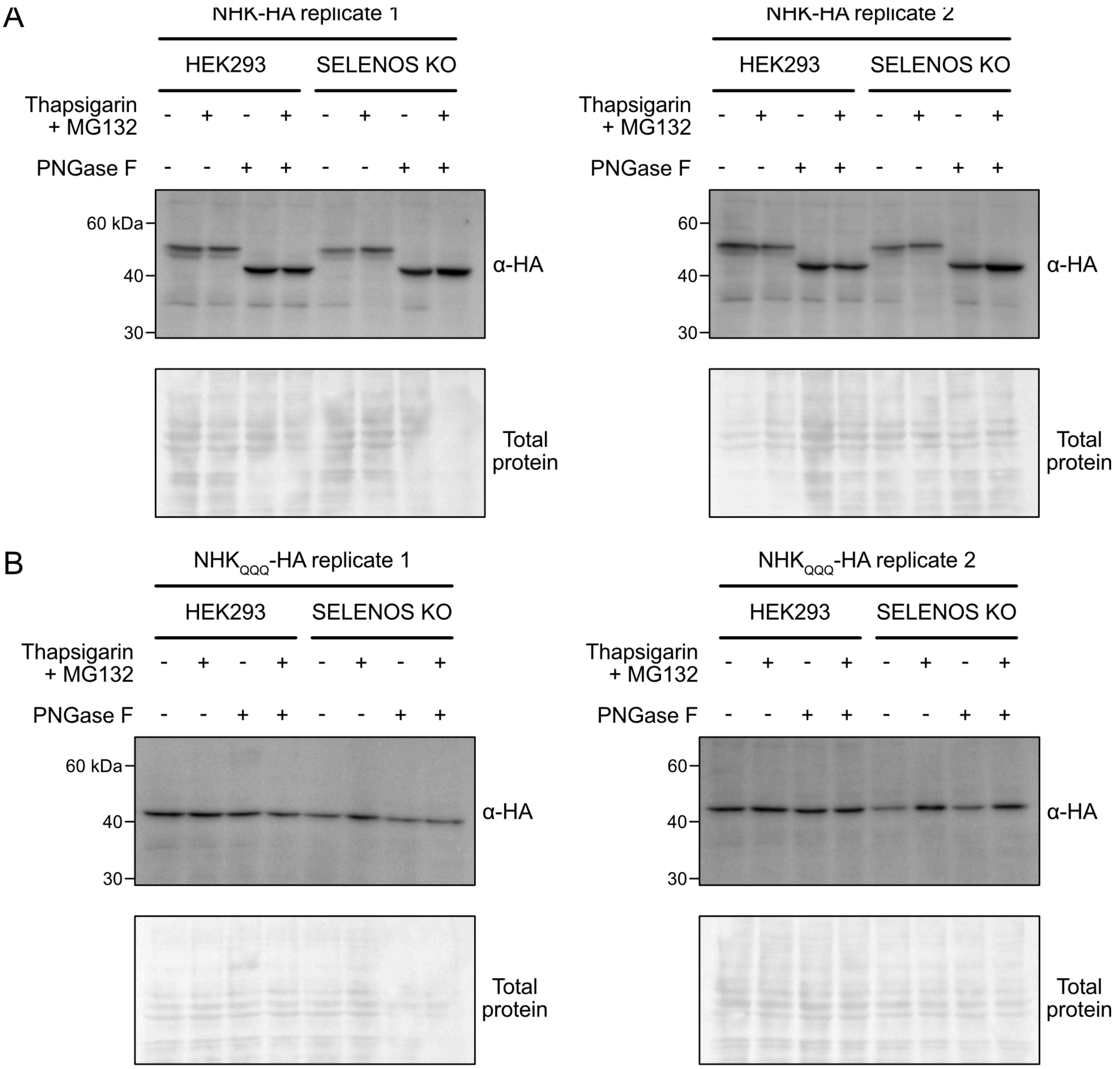


Figure S5. Selenos is not required for preemptive quality control or glycosylation of selected proteins. A) The level of glycosylated $\alpha$1-antitrypsin variant New Hong Kong (NHK, C-terminal HA tag) was not significantly altered in SELENOS knockout (KO) cells. PNGase F (20 ng/µL) was added, and the samples were incubated for 20 min at 37 °C. Loss of the glycosylated band following PNGase F treatment confirms the identity of the glycosylated species detected in the immunoblots. Total protein staining was performed using Ponceau S. B) Glycosylation-deficient New Hong Kong variant, N271Q, N107Q, N70Q (NHK$_{QQQ}$ right, C-terminal HA tag). Total protein staining was performed using Ponceau S.

| Complex/subunit | Function |
| --- | --- |
| SEC61 (Sec61 translocon) | General entry gateway for precursors with signal peptides or TMHs into the ER membrane; cooperates with TRAP, EMC, and the multipass translocon. |
| SEC61A1 | Core channel enabling ER entry of precursors. |
| SEC61B | Accessory protein |
| SEC61G | Accessory protein |
| Sec62/Sec63 (auxiliary to Sec61) | Sec62/Sec63 supports Sec61 during translocation/insertion (auxiliary factors). |
| SEC62 | Regulates gating of the Sec61 translocon and interacts with ribosomes and nascent polypeptides during insertion. |
| SEC63 | Facilitates ribosome docking and recruits BiP to drive ATP-dependent protein translocation into the ER lumen. |
| Multipass translocon (MPT) | PAT–GEL–BOS supercomplex associates with Sec61 to form the multipass translocon (MPT) and can also cooperate with EMC; BOS binding to TRAP yields a multipass-TRAP translocon. |
| NCLN | |
| NOMO1 | |
| CCDC47 | Simultaneously binds to Sec61 and ribosomes |
| TMCO1 | GEL subunit; Oxa1-related insertase component. |
| SRP (signal recognition particle) | Recognizes ER-targeting signals on nascent chains and delivers RNCs to the ER. |
| SRP19 | |
| SRP54 | |
| SRP68 | |
| SRP72 | |
| TRAP | Sec61 interaction partner; assists ER import of certain precursors; part of defined translocon architectures. |
| SSR1 | |
| SSR4 | |
| Signal Peptidase Complex (SPC) | Removes cleavable signal peptides after ER entry. |
| SPCS1 | |
| SPCS2 | |
| SPCS3 | |
| OST (oligosaccharyltransferase) | Catalyzes N-linked glycosylation of nascent proteins in the ER. |
| STT3A | The catalytic core of the OST complex |
| STT3B | The catalytic core of the OST complex |
| DAD1 | |
| DDOST | |
| MAGT1 | |
| OSTC | |
| RPN1 | |
| RPN2 | |
| TUSC3 | |
| EMC (ER membrane protein complex) | Decameric membrane protein insertase; cooperates with Sec61 and with the multipass translocon. |
| EMC1 | Provides a scaffold for the complex |
| EMC2 | A soluble cytoplasmic EMC subunit involved in guiding transmembrane domains into the membrane. |
| EMC3 | Oxa1-family insertase subunit. |
| EMC4 | |
| MMGT1 | |
| EMC6 | |
| EMC7 | |
| EMC8 | |
| EMC9 | |
| EMC10 | |
| SND pathway | An SRP-independent targeting route |
| TMEM109 | |
| BRI3BP | |
| Transmembrane helix dislocase | Removal of mislocated mitochondrial proteins |
| ATP13A1 | An ATPase that removes mistargeted mitochondrial membrane proteins from the ER. |

Figure S6. Summary of the ER protein biogenesis machinery of the components found in selenos's interactome in this study.

A

| E3 | Substrate recognition / complex organization | Substrate dislocation | E2 |
|---|---|---|---|
| SYVN1 | OS9, ERLEC1, SEL1L, EDEM1-3, DNAJC10, HSPA5, FAM8A1, HERPUD1, HERPUD2, FAF2, AUP1, SELENOS | DERL2, DERL3, DERL1(?) | UBE2J1, UBE2G2 |
| TMEM129 | ND | DERL1, DERL2, SELENOS | UBE2J2, UBE2K |
| AMFR | INSIG1, INSIG2, LMBR1L, UBAC2 | DERL1 | UBE2G2 |
| MARCHF6 | ND | ND | UBE2J2, UBE2G2 |
| RNF5 | ND | DERL1 | UBE2D3, UBE2J1 |
| CGRRF1 | ERLIN1, FAF2, UBXN4, UBXN6 | TMUB2, DERL3 | UBE2J2, UBE2N, UBE2K |
| RNF185 | TMEM259 | TMUB1, TMUB2 | UBE2K, UBE2D3 |
| RNF139 | DERL1, HM13, AUP1 | DERL1(?) | UBE2J2, UBE2G2 |
| RNF145 | INSIG1, INSIG2, FAF2 | ND | UBE2G2 |
| RNF26 | TMEM33, TMEM43, ENDOD1, TMED1 | ND | UBE2J1 |
| RNF170 | ERLIN1, ERLIN2 | ND | ND |
| TRIM13 | ND | ND | ND |

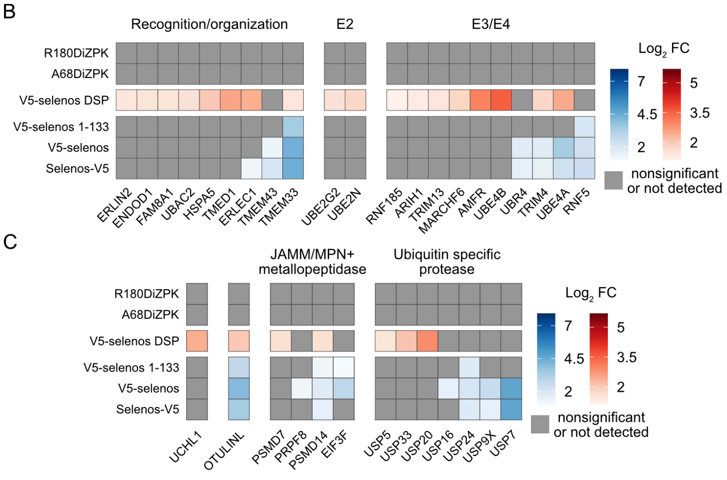


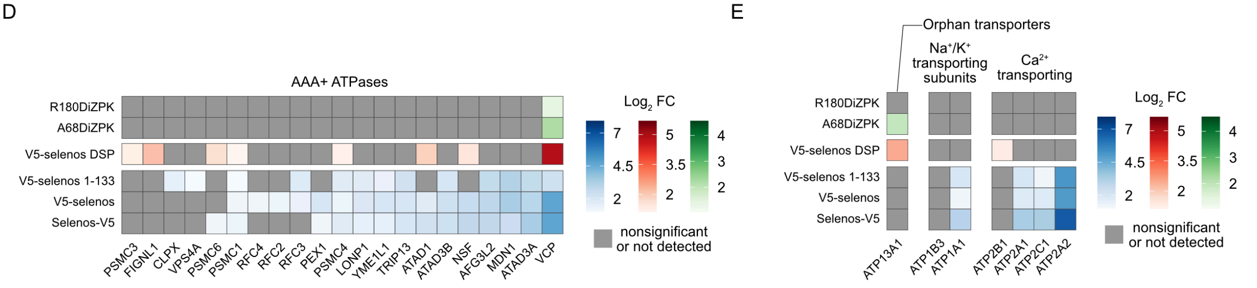


Figure S7. ERAD components and ATPases in selenos interactomes. A) The composition of ERAD assemblies for which at least one component is present in the selenos interactome. The table is adapted from reference[3]. Highlighted in red are proteins found in the selenos interactome. B-E) The $\text{Log}_2$-transformed fold change ($\text{Log}_2$ FC) from the NC-interactome experiments is shown in blue, from LC-interactomes in red, and from SC-interactome in green. Proteins that are nonsignificant or not detected under a given condition are shown in gray. Significant proteins were defined as those with an adjusted p-value below 0.05 and $\text{Log}_2$ FC above 1 (Tables S1-3). B) E2, E3 and E4 ubiquitin ligases and proteins involved in recognition/organization identified in this study that are components of ERAD. C) Deubiquitinases enriched in selenos interactome. D) AAA+ ATPases enriched in selenos interactome. E) P-type ATPases enriched in selenos interactome.

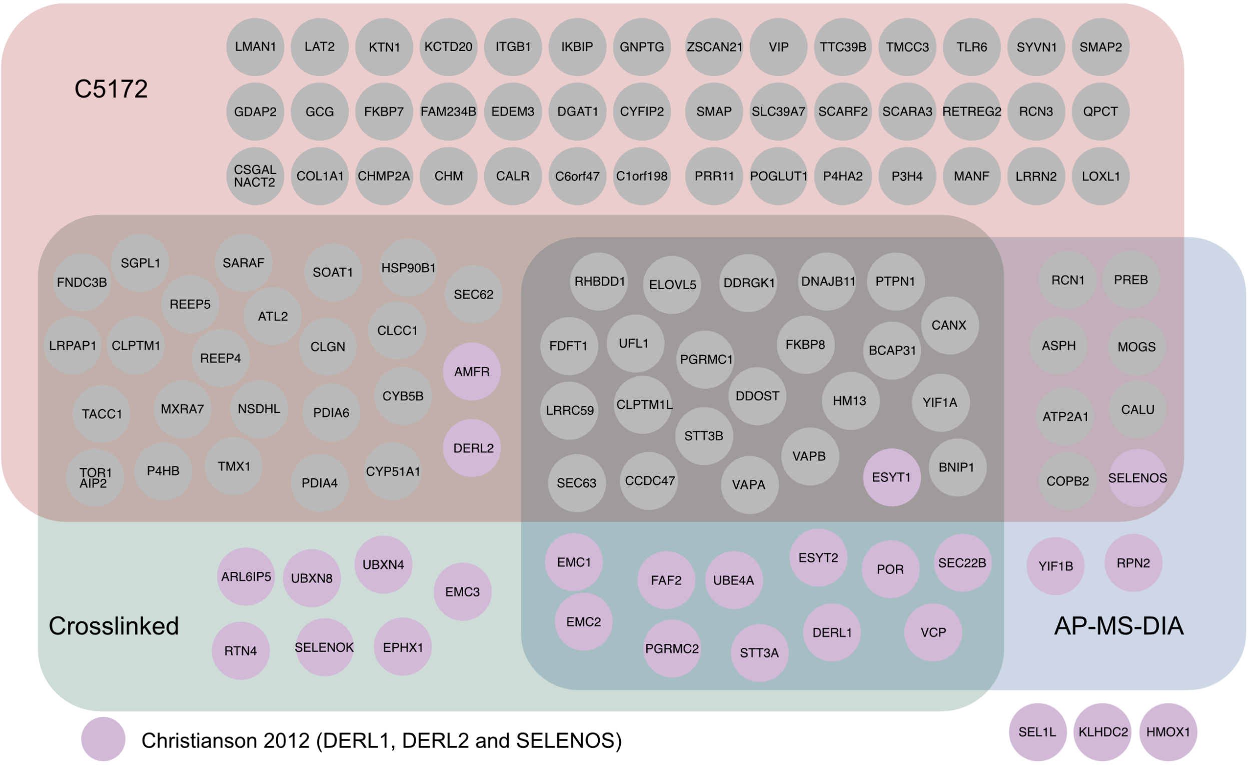


Figure S8. The overlap between selenos and derlins interactomes. The overlap between the interactome of selenos and the multimodal map of human protein assembly C5172[4] in the ER, the derlin-1 and derlin-2 interactomes published by Christianson et al.[5], and selenos interactomes reported here. “Crosslinked” refers to proteins enriched in either the SC- or LC-interactomes.

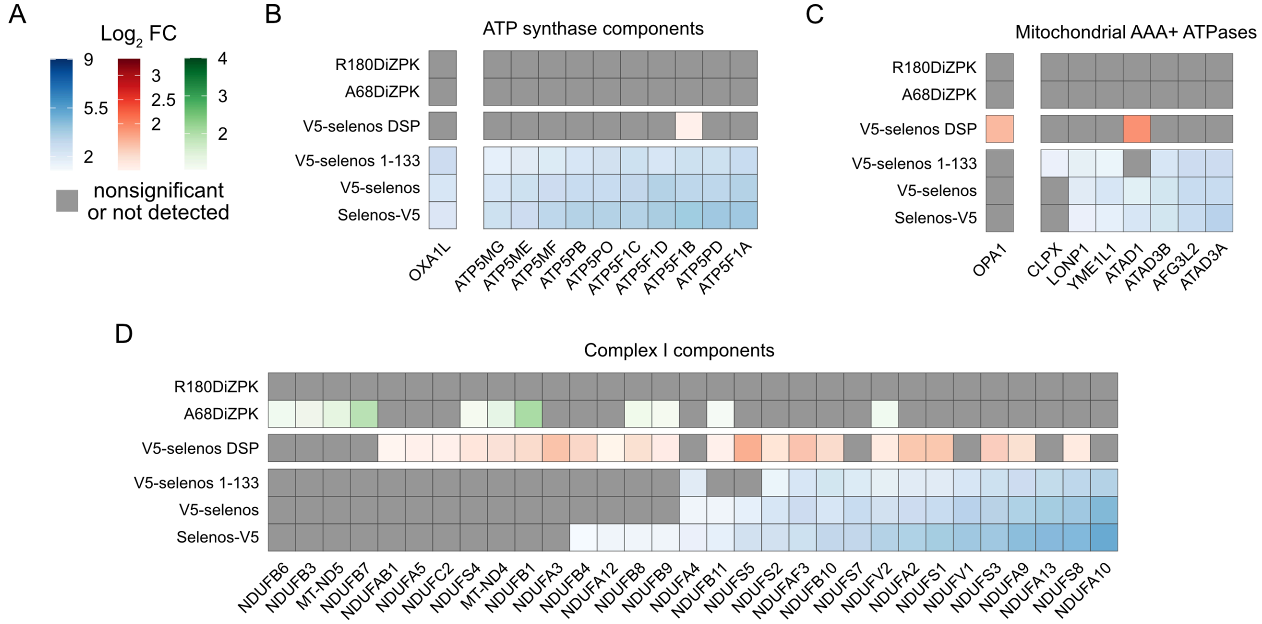


Figure S9. Mitochondrial proteins in the selenos interactome. A) The $Log_2$-transformed fold change ($Log_2$ FC) for all panels: $Log_2$ FC from the NC-interactome experiments is shown in blue, from LC-interactomes in red, and from SC-interactome in green. Proteins that are nonsignificant or not detected under a given condition are shown in gray. B-D) Significant proteins were defined as those with an adjusted p-value below 0.05 and $Log_2$ FC above 1 (Tables S1-3). B) Components of the inner membrane respiratory complex V (ATP synthase) and associated OXA1L. D) Components of inner membrane complex I (NADH:ubiquinone oxidoreductase).

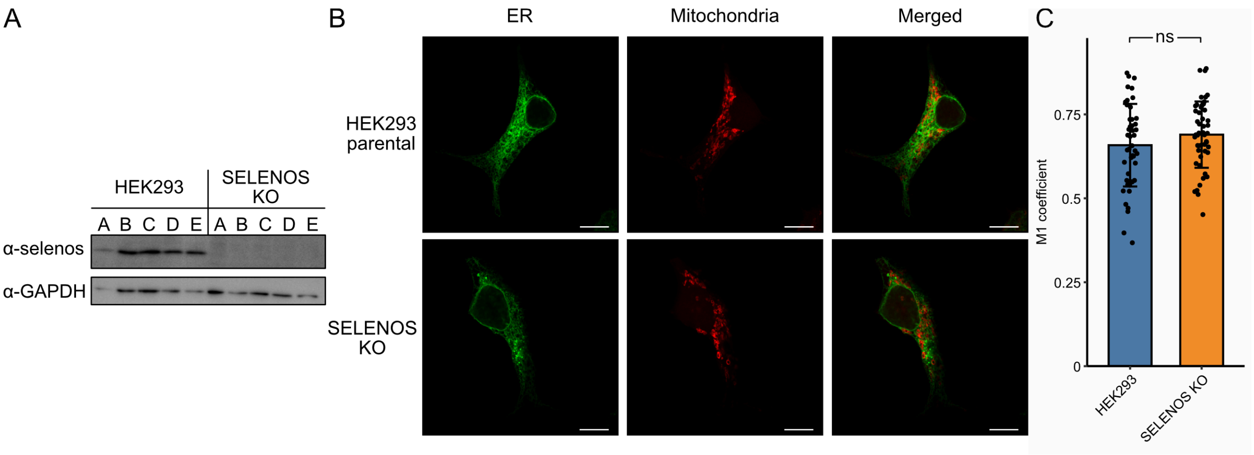


Figure S10. ER-mitochondria contact sites are not changed in SELENOS knockout (KO) and HEK293 cells. A) Immunoblot with anti-selenos antibody confirms selenos depletion in KO cells. B) Immunofluorescence microscopy of mEmerald-SEC63B (green) as an ER marker and mCherry-mito-7 (red) as a mitochondrial marker in HEK293 parental and SELENOS KO cells. C) Error bars show mean ± SD. Scale bar, 10 μm. The M1 coefficient quantifies mitochondrial localization in the ER. At least 50 images from three biological replicates were analyzed. T-test: ns, not significant.

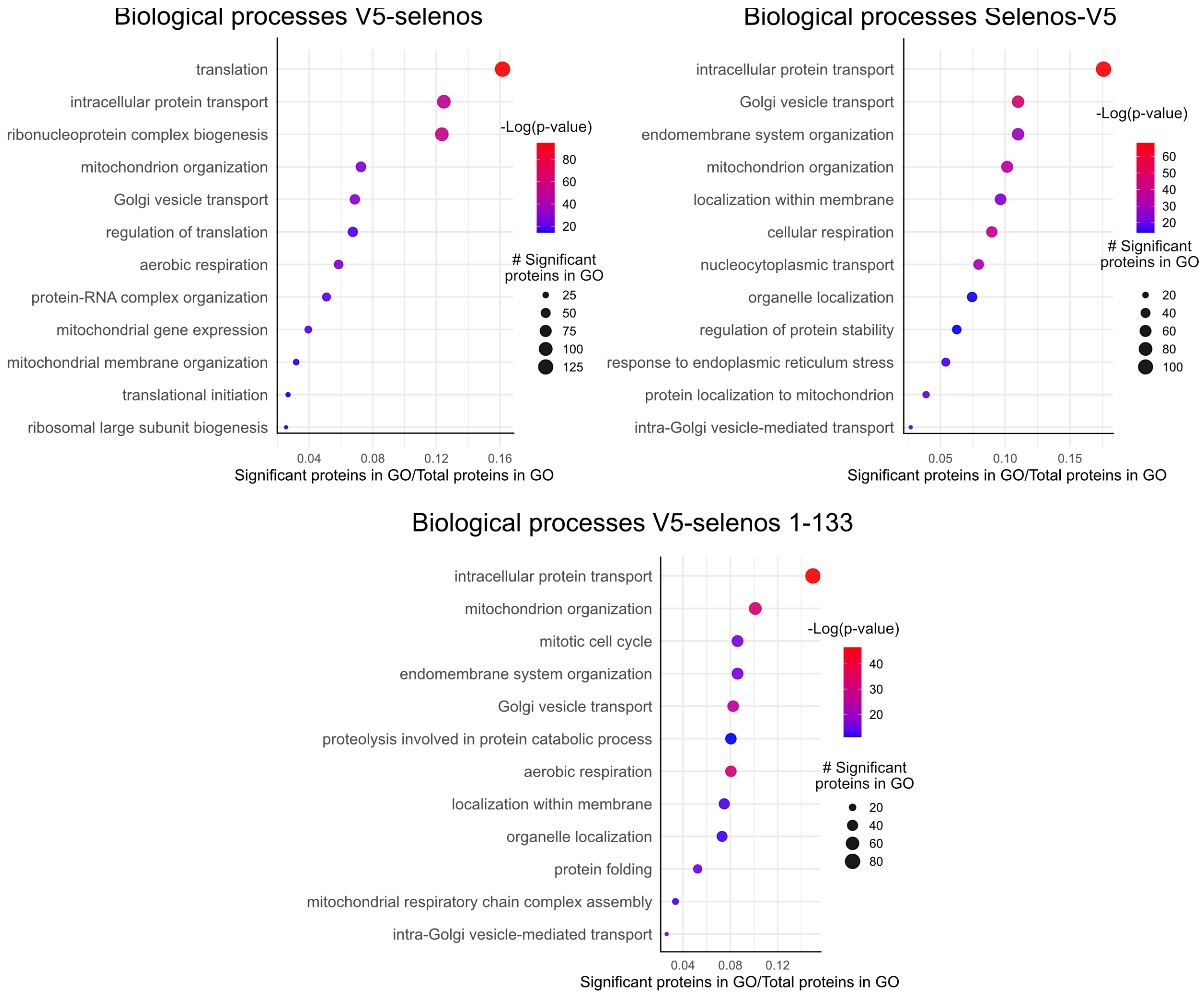


Figure S11. Gene Ontology (GO) enrichment analysis of biological processes by Metascape[6] for V5-selenos, selenos-V5 and V5-selenos 1-133. The top 12 groups (based on p-value) for each dataset are shown.

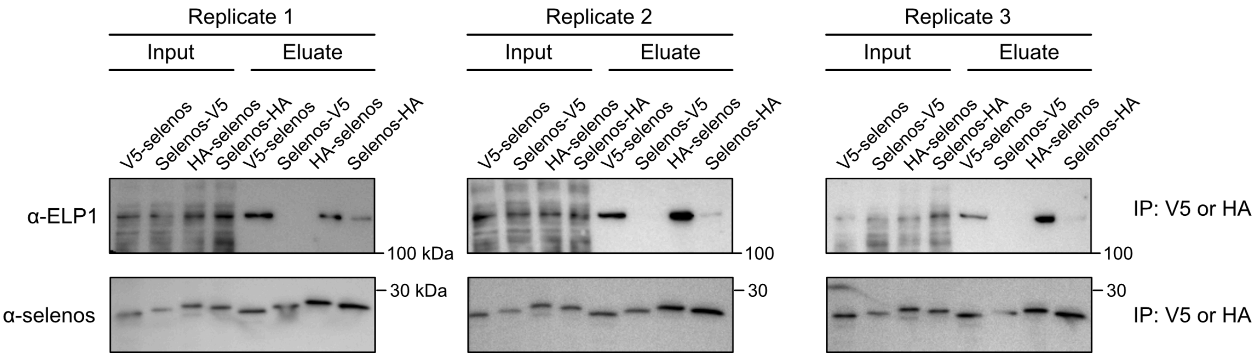


Figure S12. ELP1 levels in pulldowns are reduced when the affinity tag is positioned at the C-terminus, regardless of tag identity. Experimental procedures were identical to those described for Figure 5D. Selenos was stably expressed in Flp-In T-Rex 293 cells with either HA or V5 tags at the N- or C-terminus. HA-tagged selenos pull-downs were performed using HA-Trap magnetic beads (Proteintech, #ATMA). V5-tagged selenos was pulled down using V5-Trap magnetic beads (Proteintech, #V5TMA).

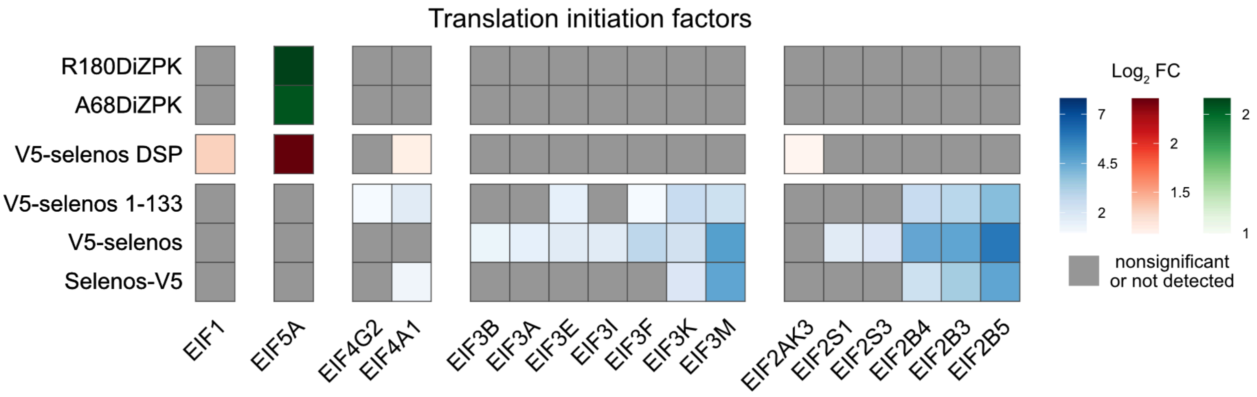


Figure S13. Translation initiation factors identified in the selenos interactomes. The Log$_2$-transformed fold change (Log$_2$ FC) from the NC-interactome experiments is shown in blue, from LC-interactomes in red, and from SC-interactome in green. Proteins that are nonsignificant or not detected under a given condition are shown in gray. Significant proteins were defined as those with an adjusted p-value below 0.05 and Log$_2$ FC above 1 (Tables S1-3).

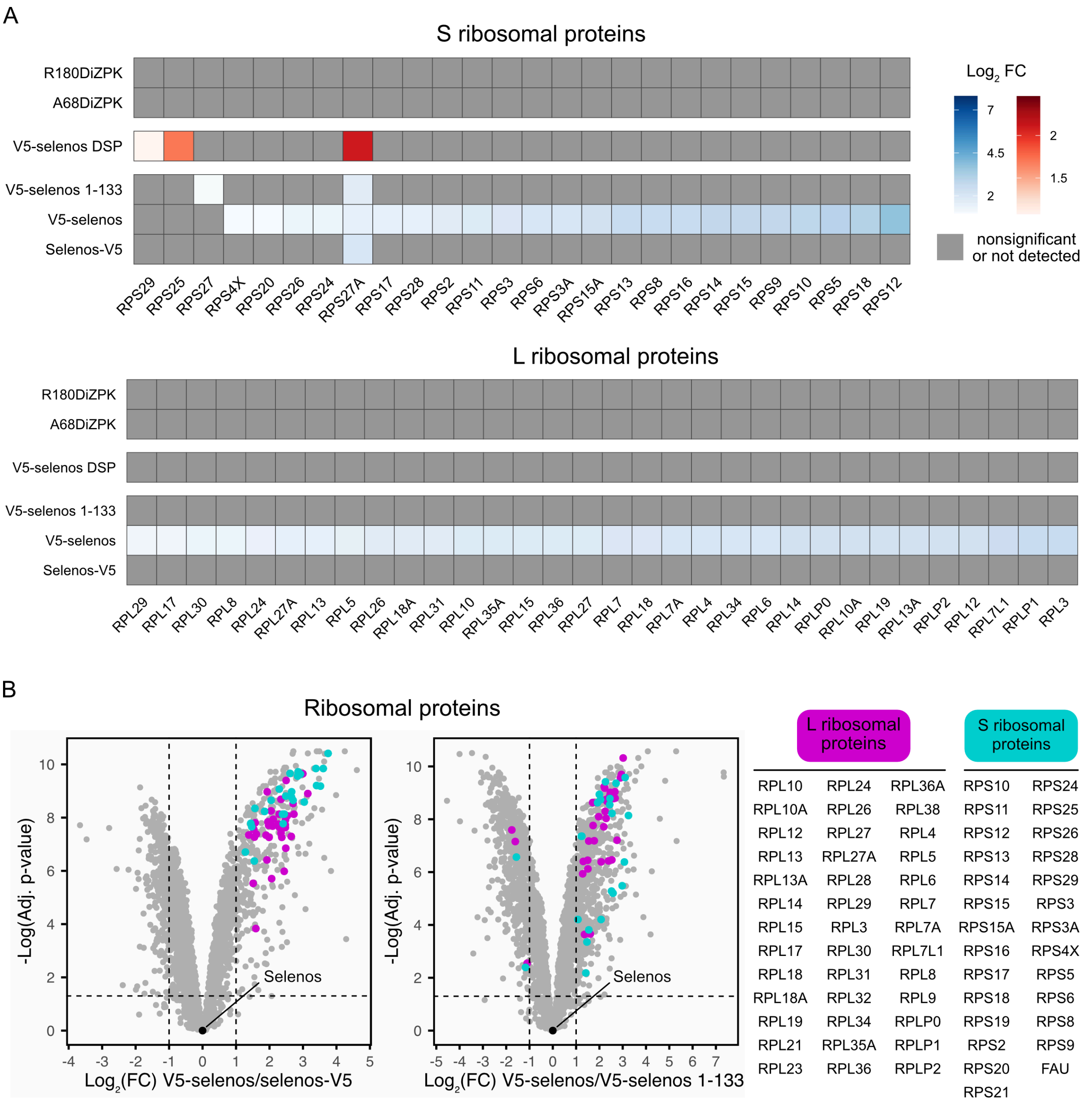


Figure S14. Ribosomal proteins identified in the selenos interactome. A) The $Log_2$-transformed fold change ($Log_2$ FC) from the NC-interactome experiments is shown in blue, from LC-interactomes in red, and from SC-interactome in green. Proteins that are nonsignificant or not detected under a given condition are shown in gray. Significant proteins were defined as those with an adjusted p-value below 0.05 and $Log_2$ FC above 1 (Tables S1-3). Ribosomal proteins of the large and small subunits are annotated according to the HGNC database. Although the overall trend is a reduction of proteins related to ribosomes when the C-terminus is not accessible, there are two exceptions to the rule that may have a functional role. The ubiquitin-ribosomal protein eS31 fusion protein (RPS27A) is moderately enriched in all conditions. In DSP-V5-selenos, the ubiquitin-ribosomal protein eL40 fusion protein (UBA52) is present. Both proteins are a source of ubiquitin but also have functions in regulating translation. UBA52 was previously shown to interact with selenos[7]. Its knockout in cells led to decreased protein synthesis and cell-cycle arrest[8]. B) When the V5 tag is placed on the C-terminus (selenos-V5), the abundance of ribosomal proteins is significantly lower compared to N-terminally-tagged selenos (V5-selenos) (Table S5). A similar reduction is observed for the truncated variant V5-selenos 1-133 relative to V5-selenos (Table S4). Proteins of the large ribosomal subunit are shown in magenta, whereas proteins of the small subunit are shown in cyan.

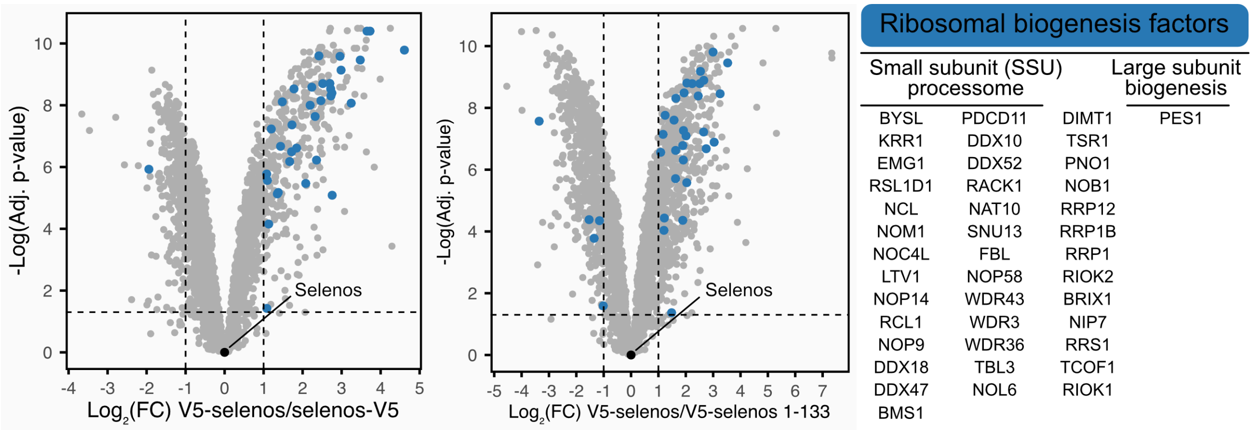


Figure S15. Ribosomal biogenesis factors identified by AP-MS-DIA. Ribosomal biogenesis factors are enriched in V5-selenos compared to selenos-V5 and V5-selenos 1-133 (Tables S4 and S5). Biogenesis factors (taken from the HGCN database) are highlighted in blue.

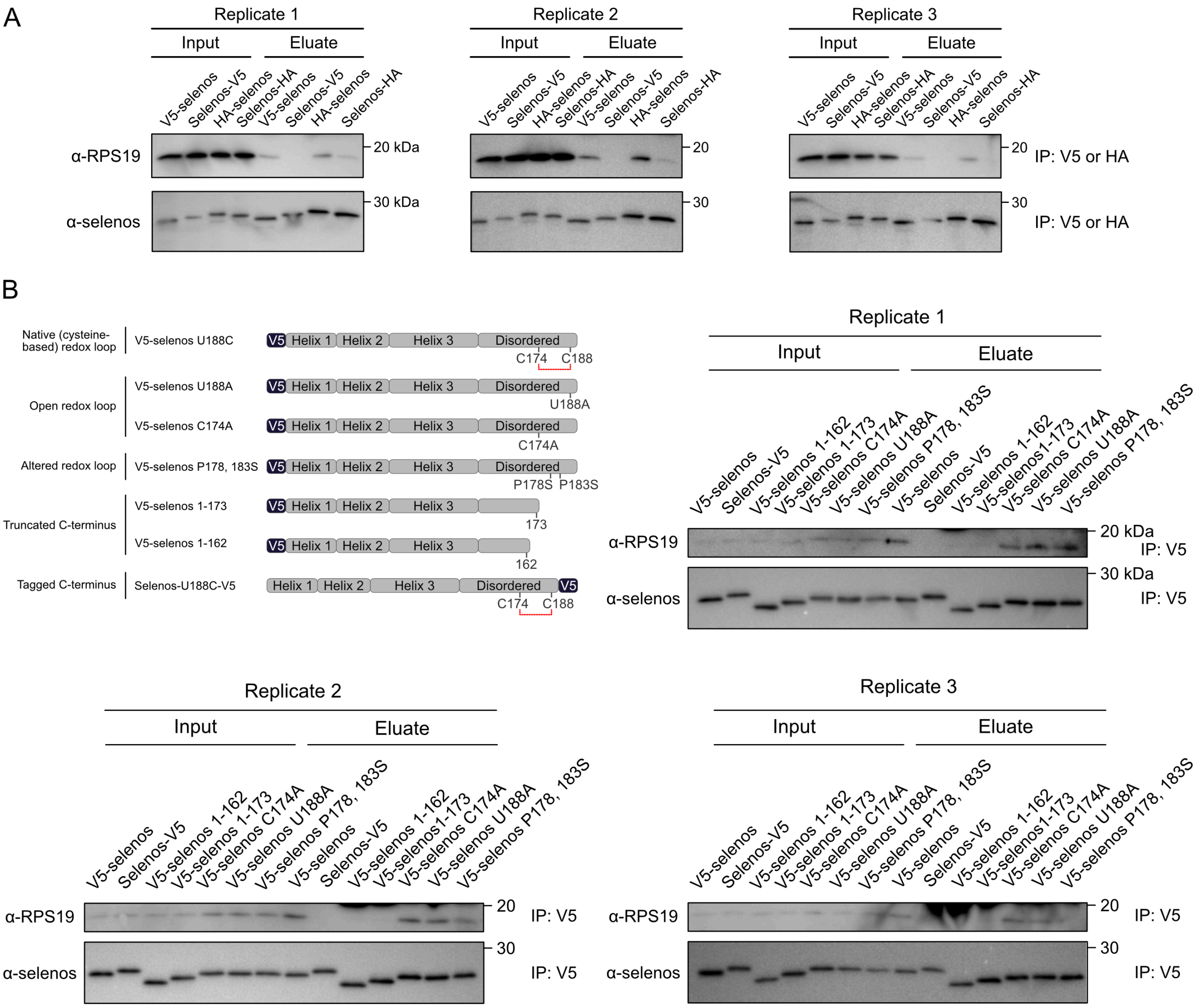


Figure S16. Selenos associates with ribosomes. (A) A C-terminal affinity tag reduces the amount of ribosomes associated with selenos, regardless of tag identity. Selenos was stably expressed in Flp-In T-REx 293 cells with either HA or V5 tags at the N- or C-terminus. HA-tagged selenos pull-downs were performed using HA-Trap magnetic beads (Proteintech, #ATMA). V5-tagged selenos was pulled down using V5-Trap magnetic beads (Proteintech, #V5TMA). Experimental procedures were identical to those described for the western blots shown in the main text. This experiment is the same as that shown in Fig. S12 but was probed for RPS19. Therefore, the selenos western blots are shared between the two figures. B) Immunoprecipitation experiments using C-terminal selenos variants confirm the AP-MS-DIA observations regarding ribosomal interactions. An antibody against RPS19 was used as a ribosomal marker. RPS19 is present in the eluates of V5-selenos but not in those of selenos-V5. Variants lacking the C-terminal redox loop show markedly reduced interaction with RPS19. Nonspecific bands close to RPS19 bands belong to bait (selenos).

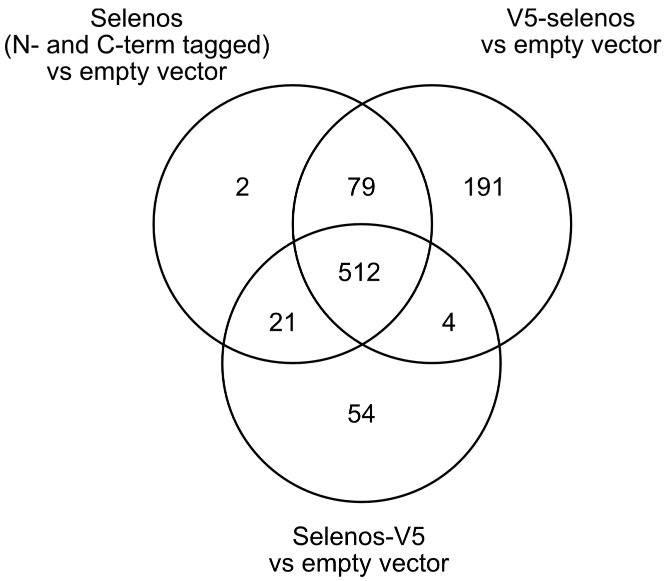


Figure S17. Grouped versus individual NC-interactome analyses of V5-selenos and selenos-V5. Comparison of the grouped NC-interactome analysis of V5-selenos and selenos-V5 versus empty-vector controls with the corresponding individual analyses of each construct versus empty-vector controls. For the grouped analysis, proteins were first subjected to replicate filtering, requiring detection in at least 4 of 5 replicates. The N-terminally and C-terminally tagged full-length selenos samples were then combined into a single group and analyzed against empty-vector controls. Proteins with at least a 2-fold increase in abundance and an adjusted P value below 0.05 were considered significant (Table S6). The grouped full-length selenos analysis was compared with the significant proteins identified in the individual NC-interactome analyses of V5-selenos and selenos-V5. A total of 512 proteins were shared between the grouped analysis and the individual analyses. The detection of these proteins in both individual datasets and in the grouped analysis supports their classification as high-confidence selenos interactors whose enrichment is independent of affinity-tag placement. This list was generally consistent with the overlap of significant proteins between the individual V5-selenos and selenos-V5 analyses, with only two additional proteins becoming significant in the grouped analysis and four proteins no longer retained. Among the 270 proteins that were significant only in the V5-selenos dataset and not in the selenos-V5 dataset, 79 were also significant in the grouped analysis. Among the 75 proteins that were significant only in the selenos-V5 dataset and not in the V5-selenos dataset, 21 were also significant in the grouped analysis.

Consistent with the individual analyses (Table S1 and Fig. S14A) and with the direct comparison of V5-selenos and selenos-V5 (Table S5 and Fig. S14B), most ribosomal proteins were exclusive to the V5-selenos dataset, confirming that C-terminal tag placement disrupts ribosomal interactions. In contrast, ELP1, ELP2, and ELP3 remained significant in the grouped analysis (Table S6). Notably, these elongator complex components were also significant in the selenos-V5 dataset, although their enrichment was lower than in V5-selenos (Fig. 5B).

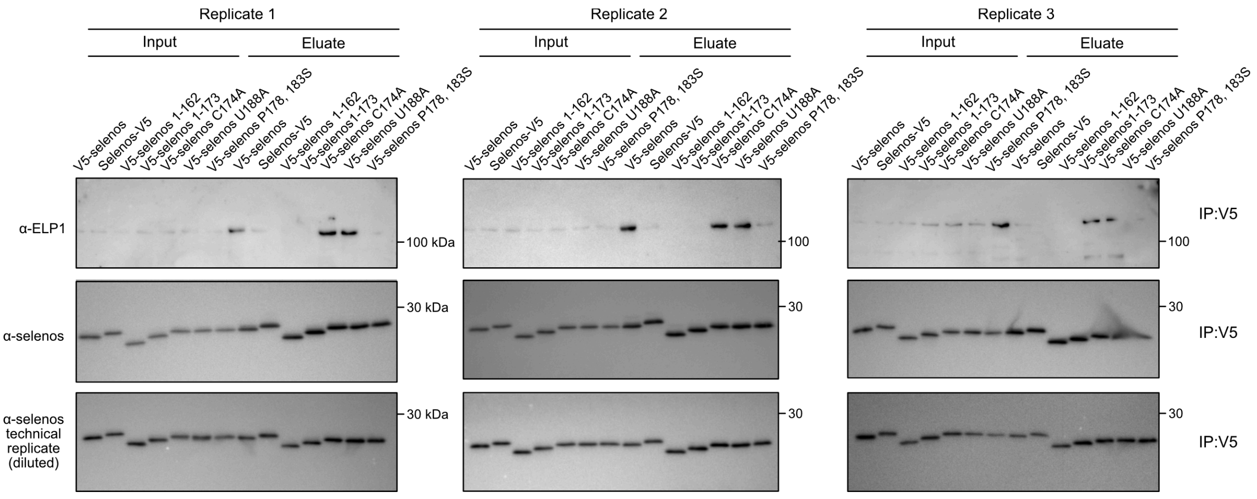


Figure S18. Biological replicates of the experiments shown in Fig. 5D and S16B.

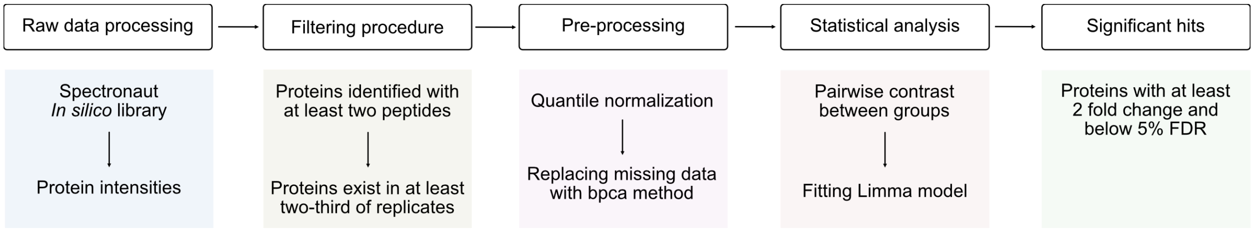


Figure S19. Workflow of the statistical analysis applied to the MS datasets.

**Supporting material and methods:**

### Immunocytochemistry and colocalization analysis

100,000 HEK293 cells or SELENOS knockout (KO) cells were seeded onto gelatin-coated coverslips (no. 1.5 thickness) in 6-well plates. HEK293 SELENOS KO cells were generated with CRISPR-Cas9 by the Gene Editing Institute at Christiana Care Hospital. Cells were grown in DMEM containing 10% (v/v) FBS and 1% (v/v) penicillin-streptomycin supplemented with 60 nM $Na_2SeO_3$. Cells were incubated in a humidified incubator at 37 °C, 5% $CO_2$, and grown to 50% confluency. Transfection was conducted using a mixture of mEmerald-SEC61B (#90992) and mCherry-mito (Addgene #55102) with Polyjet transfection reagent according to the manufacturer’s instructions and incubated for 18 h before staining.

Staining followed a modified version of the protocol described in reference[9]. Cells were rinsed three times with PBS before being fixed with 4% (w/v) formaldehyde in growth medium at room temperature for 15 min. After four PBS (137 mM NaCl, 2.7 mM KCl, 10 mM $Na_2HPO_4$, and 1.8 mM $KH_2PO_4$) washes of 5 min each, coverslips were mounted onto slides with ProLong Diamond Antifade Mountant (Invitrogen #P36961). Samples were allowed to be fixed overnight, and imaging was performed with a 63x/1.4NA/oil-immersion objective on a Zeiss LSM 880 with Airyscan. Image processing was conducted using ZEN Blue software (Zeiss).

Confocal images were imported into Imaris (Oxford Instruments), version 10.1, for quantitative analysis. Segmentation of mEmerald and mCherry channels was performed using the machine learning training functionality. For each channel, the model was trained using representative examples of foreground (true signal) and background regions. Surface objects were therefore built around structures of interest. Masked channels were then generated from the surfaces, retaining the original pixel intensities within the corresponding surface while setting all values outside the surface to zero. For colocalization analysis, overlapping channels were generated and analyzed using the “Coloc” module in Imaris. Manders’ overlap coefficients (M1) were exported, representing the fraction of intensity in one channel overlapping with the other. At least 50 images from three biological replicates were analyzed for the calculation of an unpaired two-sample Student's t-test.
